# Multivariate divergence in wild microbes: no evidence for evolution along a genetic line of least resistance

**DOI:** 10.1101/2022.09.06.506840

**Authors:** Emile Gluck-Thaler, Muhammad Shaikh, Corlett W. Wood

## Abstract

Trait evolution depends both on the direct fitness effects of specific traits and on indirect selection arising from genetically correlated traits. Although well established in plants and animals, the role of trait correlations in microbial evolution remains a major open question. Here, we tested whether genetic correlations in a suite of metabolic traits are conserved between two sister lineages of fungal endophytes, and whether phenotypic divergence between lineages occurred in the direction of the multivariate trait combination containing the most genetic variance within lineages i.e., the genetic lines of least resistance. We found that while one lineage grew faster across nearly all substrates, lineages differed in their mean response to specific substrates and in their overall multivariate metabolic trait means. The structure of the genetic variance-covariance (G) matrix was conserved between lineages, yet to our surprise, divergence in metabolic phenotypes between lineages was nearly orthogonal to the major axis of genetic variation within lineages, indicating that divergence did not occur along the genetic lines of least resistance. Our findings suggest that the evolutionary genetics of trait correlations in microorganisms warrant further research, and highlight the extensive functional variation that exists at very fine taxonomic scales in host-associated microbial communities.

## Introduction

Organisms are composed of complex assemblages of correlated traits (Lande and Arnold 1983). Consequently, the evolution of any given trait depends not only on its direct effect on fitness, but also on the indirect effects of the traits with which it is genetically correlated (Wagner 2011). Decades of research in macroorganisms has established that the strength and direction of trait correlations can either constrain or facilitate evolution, depending on their alignment with or opposition to selection, and thereby play a major role in adaptation (Lande 1979; Lande and Arnold 1983; Schluter 1996; Agrawal 2020). In contrast to macroorganisms, it remains unclear whether trait correlations are similarly important in the evolution of microorganisms like bacteria and fungi.

In plants and animals, there is extensive empirical evidence that trait correlations shape evolutionary trajectories within species (Via 1984; Strauss and Irwin 2004; Ingleby et al. 2014; Gompert et al. 2015; McGlothlin et al. 2018; Cope et al. 2021). In short-term selection experiments, this can manifest as a difference between the direction of selection and the direction of the evolutionary response. Over longer evolutionary time, this manifests as divergence among lineages that is aligned with the multivariate trait combination containing the most genetic variance (i.e., the major axis of genetic variation G_max_) (Schluter 1996). Schluter famously described this pattern as divergence along the “genetic lines of least resistance” (Schluter 1996). Substantial attention has also focused on the evolutionary lability of trait correlations: just like trait means, trait correlations can evolve via the same four evolutionary forces of natural selection, mutation, gene flow, and genetic drift and their expression is environment dependent (Lande and Arnold 1983; Shaw et al. 1995; Estes and Phillips 2006; Wood and Brodie 2015). There are empirical examples of rapid divergence in trait correlations from artificial selection experiments (Shaw et al. 1995; Steven et al. 2020), as well as between populations (Wood and Brodie 2015). Yet in macroorganisms, the dominant pattern is that trait correlations are surprisingly stable over evolutionary time (Arnold et al. 2008; McGlothlin et al. 2018, 2022).

In microorganisms, however, the evolutionary lability of trait correlations, as well as their impact on the trajectory of evolution remain major open questions. In contrast to macrobes, where several decades of research have investigated evolution through explicitly multivariate frameworks (Lande 1979; Via 1984; Ingleby et al. 2014), a multivariate perspective on trait evolution is much rarer in microbes such as bacteria and fungi (Obeng et al. 2021). There are two major reasons that trait correlations may play a different role in microbial evolution, both related to differences in the modes of inheritance that are common in microbes and macrobes. First, asexual reproduction is more prevalent in microbes than in macrobes. Although relatively few quantitative genetic studies have been performed in asexually reproducing species, existing evidence suggests that genetic covariances differ between sexual and asexual lineages (Nespolo et al. 2008).

Second, horizontal gene transfer is more common in microorganisms like bacteria and fungi (Fitzpatrick 2012; Gluck-Thaler and Slot 2015; Szöllősi et al. 2015; Arnold et al. 2022; Gluck-Thaler et al. 2022; Bucknell and McDonald 2023). As a result, microbial phenotypes are often decoupled from their phylogenetic history such that closely related lineages may not be ecologically similar (Boon et al. 2014; Djemiel et al. 2022). Horizontal gene transfer could restructure correlations between traits if the horizontally transferred genes influence some but not all of the measured traits. Alternatively, if the horizontally transferred genes influence all measured traits, horizontal gene transfer could transport populations to regions of phenotypic space that are not aligned with the major axis of trait correlations. This scenario is plausible if the measured traits are controlled by genes that are typically located close together in microbial genomes, as is often the case for metabolic phenotypes (Wisecaver et al. 2014; Urquhart et al. 2023). Taken together, these observations suggest (1) that trait correlations may be more labile in microorganisms than in macroorganisms; and (2) that trait correlations may have a weaker impact on the trajectory of evolution in microorganisms than in macroorganisms.

The metabolic traits of host-associated microbes are an excellent system in which to test these hypotheses. Host-associated microbes spend much of their lives exposed to complex mixtures of host metabolites (Gershenzon et al. 2012). Consequently, the growth of a host-associated microbe is a multivariate trait that can be decomposed into its ability to metabolize different host substrates (Rokas et al. 2018; Slot and Gluck-Thaler 2019; Westrick et al. 2021). In plants, for example, carbohydrates such as sugars, fibers and starches are often the principal source of carbon accessible to microbes and are growth-promoting resources (Liu et al. 2011). Yet plants also produce secondary metabolites that play multifaceted roles in signalling, growth, and defense (Piasecka et al. 2015). Phenolics, for example, are important constituents of plant cell walls and critical players in constitutive and inducible defense responses against microbial pathogens (Nicholson and Hammerschmidt 1992; López-Goldar et al. 2018; Wallis and Galarneau 2020). As a result, the ability to degrade phenolic compounds is an important determinant of microbial growth on plant tissues (Schäfer et al. 1989; Kliebenstein et al. 2005; Zhao et al. 2019; Nickerson et al. 2023). The simultaneous presence of growth-inhibiting compounds (like secondary metabolites) and growth-promoting compounds (like carbohydrates) suggests that trade-offs between stress tolerance and rapid growth may be an important axis of trait variation in host-associated microorganisms.

In this study, we compared genetic correlations in a suite of metabolic traits between two lineages of endophytic fungi. We further tested whether divergence in mean metabolic phenotype between lineages occurred along the genetic lines of least resistance (sensu Schluter 1996). The endophyte lineages we used in this study are dark septate endophytes, a polyphyletic group of melanized Ascomycete fungi that dominate plant microbiomes across all major plant groups in terrestrial ecosystems (Ruotsalainen et al. 2021). Dark septate endophytes from the Molliciaceae family frequently colonize the roots of *Vaccinium* species, a widely-distributed genus of shrubs that includes blueberry and cranberry (Gorzelak et al. 2012; Yang et al. 2018; Morvan et al. 2020). Individual Mollisiaceae species and even individuals within the same species often differ dramatically in their interactions with plants, ranging from commensal to mutualistic to pathogenic (Newsham 2011; Tellenbach et al. 2011; Reininger et al. 2012), suggesting variation within and between species and lineages has important ecological consequences. Yet despite their potential to impact plant host function and their ubiquity, our understanding of phenotypic differences among dark septate endophytes and of the ecological factors shaping their distributions is still in its infancy (Tanney and Seifert 2020).

We isolated two lineages of Molliseaceae endophytes from the roots of lowbush blueberry (*Vaccinium angustifolium*) across three sites in Eastern North America. We then compared endophyte growth across a panel of phenolic and carbohydrate metabolites (to capture their response to specific host chemicals) and extracts of whole plant tissues and organs (to capture their response to more realistic biochemical environments), and tested for differences in trait means and correlations between lineages. We used these two experiments to answer three questions: (1) Have these endophyte lineages diverged in their response to each metabolite and extract, and in their multivariate mean metabolic phenotype? (2) Are there genetic trade-offs between growth on phenolics and carbohydrates, and if so, are these trade-offs conserved across lineages? and (3) Did divergence in mean metabolic phenotypes occur along a genetic line of least resistance (sensu Schluter 1996)?

## Materials and Methods

### Endophyte and plant material collection

Lowbush blueberry (*V. angustifolium*) plants were sampled from June to October 2020 from three second growth forests in the Eastern United States that are primarily used for scientific research: the research forest at Chatham University’s Eden Hall Campus in Richland Township, Pennsylvania (40°39’50.0“ N 79°57’12.3” W); Powdermill Nature Reserve in Westmoreland County, Pennsylvania (40°09’41.1“ N 79°16’28.9” W); and Mountain Lake Biological Station in Giles County, Virginia (37°22’34.2“ N 80°31’12.1” W). At each location, we selected 2-7 sites located at least 500m apart, and within each site we haphazardly sampled 5-10 *V. angustifolium* individuals located at least 10m apart with no observable blueberry plants between them. Fine fibrous roots located no deeper than 10cm from the soil surface were collected from individual plants using a trowel and knife, stored in plastic bags, and used immediately that same day for endophyte isolation.

At Mountain Lake Biological Station in October 2020, we collected additional plant materials to use in the preparation of growth media for two experiments (see below). Whole root systems and attached leaves from approximately 50 *V. angustifolium* plants growing from three large patches were collected on a single day. Root systems and leaves were pooled across sample sites. We also haphazardly collected fallen leaves from *Quercus* spp. (oak) and *Acer* spp. (maple), and fallen needles from a stand of planted *Pinus monticola* (western white pine). Intact plant leaves were collected from the ground and not from living trees to ultimately extract material that Molliseaceae endophytes could be expected to encounter during the saprophytic stage of their lifecycle around the roots of their lowbush blueberry hosts (Tanney and Seifert 2020). Since the leaves were collected from the ground, we were not confident in ID’ing them to the species level. All roots and leaves were kept at -80°C for approximately four months until used for media preparation.

### Endophyte isolation and genotyping

Endophytic fungi were isolated from *V. angustifolium* roots using a sequence of ethanol/bleach sterilization steps followed by direct plating onto 50% strength Modified Melin-Norkrans agar (MMN; Supplementary Methods). A total of 69 fungi isolated from *V. angustifolium* roots with melanized morphologies characteristic of dark septate endophytes were successfully genotyped by sequencing the internal transcribed spacer (ITS) barcoding gene (Table S1)(Gardes and Bruns 1993). We identified each isolate to taxonomic family level using a BLASTn search of their ITS sequence against the NCBI nr/nt database (last accessed 01/31/2021). The resulting sample of ITS sequences from Mollisiaceae isolates, along with reference sequences from known species within this family (Tanney and Seifert 2020), were aligned using mafft (--auto)(Katoh and Standley 2013). Pairwise sequence identities between isolates were calculated based on this alignment. Alignments were trimmed using ClipKit (default settings)(Steenwyk et al. 2020), and used to build a maximum likelihood phylogeny with IQtree (-m MFP -b 100) (Nguyen et al. 2015). This phylogeny revealed that 52 isolates could be confidently placed within the Mollisiaceae family.

Given that species-level classifications are notoriously difficult to resolve in the Mollisiaceae, we used the phylogenetic placements of our isolates relative to the closest described species to provisionally classify them into one of two lineages: the *Phialocephala fortinii* s.l. – *Acephala applanata* species complex (PAC) and “Clade D” sensu Tanney and Seifert 2020. PAC contains some of the most well-studied and frequently recovered dark septate endophyte species from coniferous and ericaceous plant roots (Grünig et al. 2008). Clade D is an ad-hoc phylogenetic group which includes *Mollisia nigrescens* and other poorly described root endophytes that have previously been recovered from *Vaccinium* tissues (Tanney and Seifert 2020).

We initially selected n_Clade D_ = 13 and n_PAC_ = 17 isolates to phenotype from our available collection by prioritizing isolates with different ITS sequences and/or that were recovered from different host individuals to minimize the phenotyping of clones (Figure 1). While both Clade D and PAC isolates were recovered from the Mountain Lake site (n_Clade D_ = 10; n_PAC_ = 14), only Clade D isolates were retained from the Chatham site (n_Clade D_ = 3) and only PAC isolates were retained from the Powdermill site (n_PAC_ = 3). We ultimately filtered out several isolates due to quality control issues, resulting in n_Clade D_ = 11 for the Metabolite Experiment, n_Clade D_ = 13 for the Extract Experiment, n_PAC_ = 15 for the Metabolite Experiment and n_PAC_ = 16 for the Extract Experiment (Supplementary Methods) (Parks and Goldman 2014). It is important to note that between 11-16 isolates is on the low end of the sample size used to estimate G-matrices. In a meta-analysis of 95 G-matrices, Wood and Brodie III (2015) found that the median sample size used to estimate a G-matrix was 38.5 families or genotypes (mean = 52.2, range = 5-587). Therefore, while our sample size is within the range of past G-matrix studies, we have relatively limited statistical power to identify small differences in G-matrix structure. In our study, the small number of isolates per lineage is a feature of the challenge of targeting specific microbes for sampling in the field: unlike most macrobes, a microbe’s identity cannot be determined without culturing or sequencing it.

**Figure 1:**
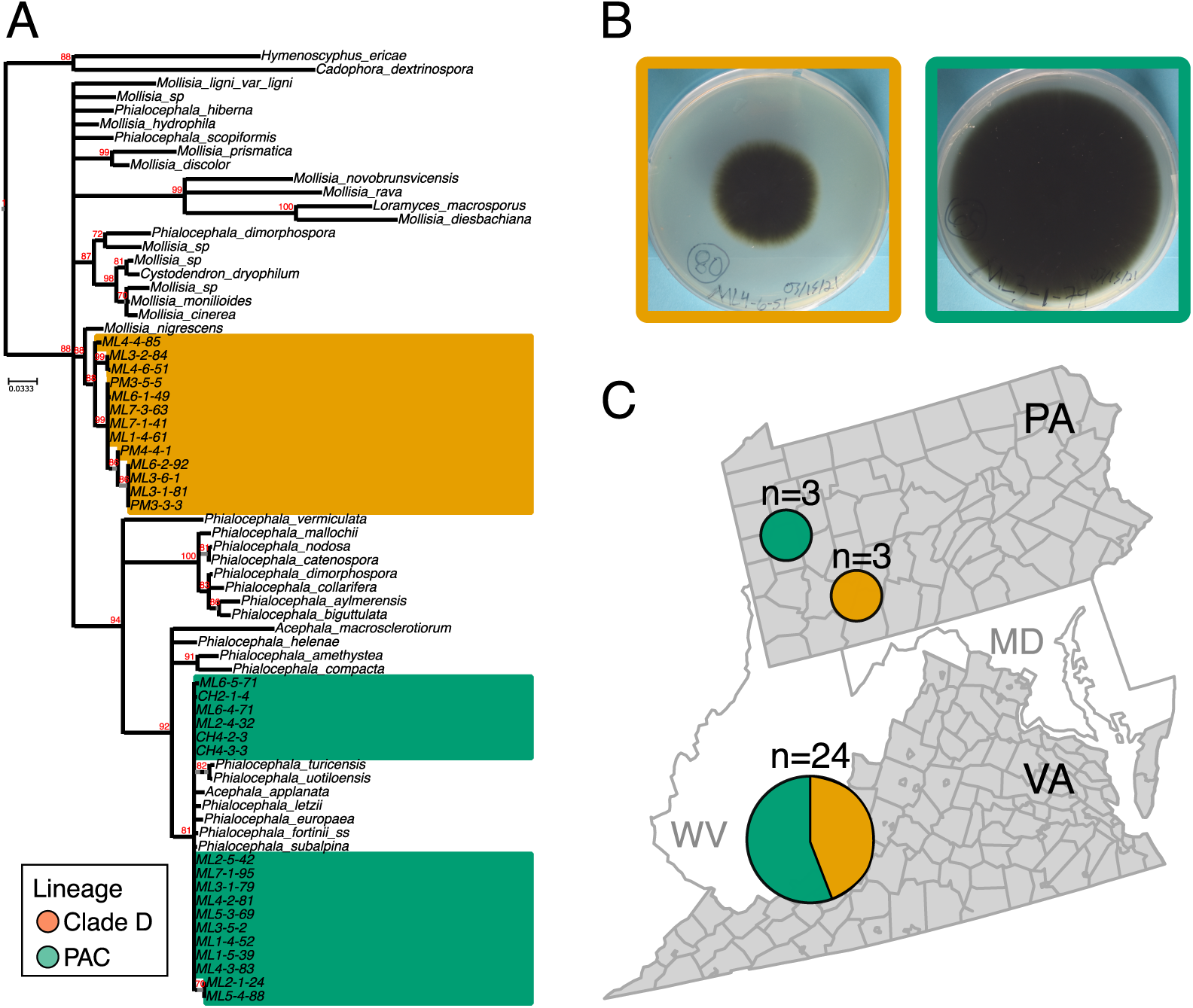
Two lineages of Mollisiacean endophytes sampled from wild *Vaccinium angustifolium* roots across multiple sites in Eastern North America. A) A maximum likelihood phylogeny of the ITS sequences of all Mollisiacean fungi isolated as part of this study and reference sequences obtained from public databases. Isolates from this study are highlighted in green if part of the *Phialocephala fortinii* s.l. – *Acephala applanata* species complex (PAC), highlighted in orange if part of the ad-hoc phylogenetic group Clade D, and reference sequences are not highlighted. Bootstrap support is labeled in red and all branches with <70% bootstrap support have been collapsed. B) Pictures of typical Clade D (left) and PAC (right) isolates after two weeks growth on 1% blueberry root extract media C) Approximate locations of sample sites in Pennsylvania (PA) and Virginia (VA), with the proportion of each lineage from each site visualized as a pie chart.

We conducted all statistical analyses described below using isolates from all three sites as well as a more restricted set containing only isolates from Mountain Lake to test whether including isolates from different geographic areas changed our results. Limiting our analysis to Mountain Lake isolates only does not qualitatively change the results of our study. Therefore, we choose to present results from the analysis of isolates from all three sites.

### Plant metabolite experiment: media preparation and experimental design

We compared metabolic trait means and metabolic trait covariances within and between lineages, where we define a metabolic trait as growth on a particular plant-derived substrate. We selected 6 phenolic and 5 carbohydrate metabolites produced across different organs in *Vaccinium* and therefore likely encountered by fungal endophytes colonizing species within this genus (Table S2). We focused on phenolics and carbohydrates to capture potential trade-offs between performance on putatively growth-inhibiting and growth-promoting compounds (however, despite the fact that phenolics are an important component of inducible defenses in plants (Nicholson and Hammerschmidt 1992; López-Goldar et al. 2018; Wallis and Galarneau 2020), they did not inhibit growth in our experiment; see Results). We added each metabolite dissolved in ethanol to autoclaved media amended with a hot water extraction of *V. angustifolium* roots (Supplementary Methods). Each of the 11 treatments consisted of known physiological concentrations of each metabolite (Stribley and Read 1974; Harris et al. 2007; Riihinen et al. 2008; Kaur et al. 2012; Edwards et al. 2018; Ștefănescu et al. 2020). We used media amended with host root extract to capture the effect that each metabolite may have on endophyte growth in the context of other host-associated metabolites, as opposed to adding metabolites to standardized minimal media. As a control treatment, we added 20ml of pure ethanol to root extract media, for a total of 12 treatments.

We ran this experiment in three cohorts separated by 1-2 weeks each due to the large number of plates. Each cohort had one replicate of each isolate-metabolite combination (34 isolates x 12 treatments = 408 plates / cohort). All plates for a given cohort were prepared on the same day using root extract from one of the six original bags of *V. angustifolium* root system collections (see above) and left to dry for 48 hours in a laminar flow hood. Plates were then inoculated with 4mm plugs from two-week old cultures grown on MMN plates, sealed with micropore tape, haphazardly arranged and incubated upside-down in the dark at room temperature.

### Plant extract experiment: media preparation and experimental design

We selected root and leaf materials from *V. angustifolium,* as well as leaves from three different plant species sympatric with *V. angustifolium* (*Pinus monticola, Quercus* spp. and *Acer* spp.). We used these additional plant species to create ecologically relevant variation in multivariate biochemical environments for the plant extract experiment. In contrast to the metabolite experiment described above, this experiment was designed to capture potential trade-offs in performance across more realistic biochemical environments. We prepared root and leaf extracts from materials collected at Mountain Lake Biological Station (see above) using a hot water extraction method (Supplementary Methods).

Autoclaved extracts were added to autoclaved DI water agar (15g/L) to a final concentration of 5g/L, and the resulting solutions were immediately poured into plates prepared and left to dry as above. Final extract concentrations in this experiment were 5 times greater than those in the metabolite experiment because they were intended as the main treatment effect. There were 3 replicates per isolate x extract treatment (34 isolates x 5 treatments x 3 replicates = 510 plates), and a single experimental cohort. All plates were prepared on the same day and inoculated and stored as above.

### Measuring growth rate

Given that many Mollisiaceae endophytes primarily reproduce through clonal vegetative growth (Tanney and Seifert 2020), growth rates are expected to be important ecological traits (Aguilar-Trigueros et al. 2015). We measured colony area in cm^2^ at 7, 14, and 21 days after inoculation and calculated the Area Under the Growth Progress Stairs (AUGPS) as a quantitative measurement of growth over time, such that AUGPS units were cm^2^ x 7 days (Supplementary Methods). We calculated AUGPS using the audps function (type = “absolute”) in the R package Agricolae (Mendiburu and Simon 2015). While all isolates were able to grow on all treatments, 30 out of the 1,530 plates prepared across all experiments had cultures that did not grow at all over the 21-day period, as occasionally happens during culture-based experiments, and were subsequently discarded. This resulted in at most 1 replicate being discarded for each genotype x treatment combination.

Haphazardly selected groups of plates were imaged using a vertically mounted Nikon D3500 camera equipped with a Nikon AF-P DX 18-55mm f/3.5-5.6G VR lens. A ruler was placed in each image and was used to calibrate measurements. Plates were illuminated from two sides by mounted fluorescent lights. We measured colony area from images of each plate using ImageJ software (Schneider et al. 2012), where the boundaries of each colony were estimated by eye. The colony area at day 0 for all isolates (0.113 cm^2^) was determined based on a random sample of 10 plates imaged at this time. Colonies without identifiable boundaries were marked as missing. Images from the extract experiment taken on day 21 were discarded because the media and fungal tissues progressively darkened over this period of time - perhaps due to increased melanization or the activity of secreted enzymes - such that the fungal colony boundaries could not reliably be identified. Growth rates for the Extract Experiment were therefore estimated using two timepoints only, in addition to the starting timepoint at day 0, which limited our ability to capture variation in growth rates between day 14 and 21.

### Statistical analysis: Linear mixed models

Prior to our analyses, we removed 4 isolates originally assigned to Clade D from our dataset due to poor phylogenetic support (Supplementary Methods). In addition, we removed 4 isolates (2 PAC and 2 Clade D) from the metabolite experiment and 1 PAC isolate from the extract experiment due to possible contamination (Supplementary Methods). We performed all statistical analyses in R 4.1.1 (R Core Team 2016) and unless specified otherwise, created data visualizations using ggplot2 (Wickham et al. 2021). We constructed three different linear mixed models to test for the effects of various predictor variables on isolate growth (i.e., AUGPS) using the “lmer” function in the lme4 package (Bates et al. 2015). We included isolate and isolate-by-treatment interactions as random effects in all models to control for isolate-specific variation.

To test how individual metabolites affected isolate growth in the two endophyte lineages, we constructed a model with metabolite, phylogenetic lineage, metabolite-by-lineage interactions and cohort as fixed effects. To test how chemical class (phenolic or carbohydrate) interacts with lineage membership, we constructed a model with chemical class, phylogenetic lineage, class-by-lineage interactions and cohort as fixed effects and individual metabolite as a random effect to control for metabolite-specific effects independent of chemical class. We fit cohort as a fixed effect rather than a random effect because our experiment included only 3 cohorts, and random-effect variance estimates are imprecise for random effects with fewer than 5-6 levels (Harrison et al. 2018). Finally, to test how plant extracts interact with lineage membership, we constructed a model that included extract treatment phylogenetic lineage and extract-by-lineage interactions as fixed effects.

We tested for the significance of fixed effects with type III sums of squares using the Anova function in the car package (Fox and Weisberg 2011). We confirmed that the residuals within each model met the required assumptions of normality and homoscedasticity by simulating model residuals using the DHARMa package (Hartig and Lohse 2021) and inspecting the resulting quantile-quantile plot and the plot of the residuals versus the predicted values to detect deviations from the expected distribution. Estimated marginal means and their 95% confidence intervals for each treatment within each lineage were calculated and compared to each other using the emmeans package and post-hoc Tukey tests (Lenth et al. 2021).

### Statistical analysis: Comparisons of multivariate G-matrices

To test for differences in the multivariate response to plant chemistry between the two lineages, we calculated and compared the genotypic variance-covariance (**G**) matrices from each lineage for the plant metabolite and plant extract experiments, in addition to carbohydrate- and phenolic-specific G-matrices. In these G-matrices, each isolate trait is its mean-centered growth (AUGPS) averaged across its replicates on a specific metabolite or plant extract. We mean centered AUGPS for each isolate by subtracting the lineage mean from its untransformed AUGPS value, in a process analogous to size correction that is used when comparing different macroorganism species (McGlothlin et al. 2018), in order to increase our power to detect differences in growth due to treatment effect and not differences due to baseline growth rates between lineages. In order to avoid artificially inflating similarity between G-matrices, we removed metabolite treatments from the metabolite experiment that had no statistically significant effect on growth in either lineage (Table S3). This resulted in the removal of two phenolic treatments (chlorogenic acid and vanillic acid) and one carbohydrate treatment (starch). Removal of these treatments do not quantitatively change the results of any multivariate statistical analyses (data not shown). For the metabolite experiment, we averaged growth across cohorts (which were equivalent to experimental replicates) and for the extract experiment, we averaged across replicates. We decomposed each G-matrix into its eigenvectors and eigenvalues using base R’s prcomp function (scale = FALSE).

We calculated three test statistics that capture differences in matrix structure, following previously established methods (Wood and Brodie III 2015): the difference in total genetic variance between endophyte lineages (ΔVt), the difference between the proportion of variance along Gmax (ΔGmax), and the difference in the orientation of Gmax (θ). Gmax represents the largest eigenvector of the covariance matrix and is the linear combination of traits that captures the largest proportion of variation in growth among isolates. A large ΔVt indicates that isolates within one lineage harbor more genetic variation than do isolates in the other lineage. A large ΔGmax indicates that growth on different substrates is more strongly correlated in one lineage than in the other. A large θ indicates that the lineages differ in the direction of trait correlations. To test whether ΔVt, ΔGmax, or θ differed between lineages, we permuted isolates among the two lineage matrices 500 times and recalculated the three test statistics as above from the permuted data to generate null distributions.

To quantify similarity in overall matrix structure, we calculated correlations between the response to random vectors applied to each lineage’s G-matrix using the RandomSkewers function from the evolqg package with 10,000 random gradients (Melo et al. 2016). The random skewers (RS) method applies random vectors of selection gradients to each matrix and measures the correlation between the response vectors of the two matrices. The responses of matrices with similar structures will be highly correlated.

To examine pairwise correlations between growth on different substrates, we constructed Spearman correlation matrices using base R’s cor function. The significance of each correlation coefficient was visualized and calculated using the ggcorplot package (Kassambara 2019). Pairwise trait correlations between lineages were compared using the cocor package using the Fisher’s z test (Diedenhofen and Musch 2015). *P* values were corrected for multiple comparisons using the Benjamini-Hochberg method implemented in base R’s p.adjust function.

To test for differences in multivariate trait means between lineages, we performed a one-way MANOVA for each experiment where the different substrates were the response variables and lineage was the predictor. Here, we also mean centered trait values by subtracting mean lineage growth across all treatments within a specific experiment from each trait value. This allowed us to remove the overall growth difference between the lineages while preserving the differences between treatments within each lineage. We evaluated the effect size of lineage on substrate using the package effectsize (Ben-Shachar et al. 2020).

We also used mean centered AUGPS values to calculate Schluter’s theta according to the procedure described in Schluter (1996) so we could focus on differences between lineages due to substrate treatment and not basal growth rate (e.g., analogous to using size-corrected species means for animals (McGlothlin et al. 2018)). First, we extracted Gmax from each lineage’s G-matrix for each experiment using the prcomp() function in R. Then, we calculated the vector between each lineage’s multivariate mean (z), and normalized z such that z’z = 1. Finally, for each lineage, we calculated the angle theta in degrees between its Gmax and z.

## Results

### Lineages differ in baseline growth rates and in their growth on individual plant metabolites, chemical classes and plant extracts

For the plant metabolite experiment, we found that lineages differed in their baseline growth (lineage effect: χ^2^_df=1_ = 175.43, *P* < 0.0001), that metabolites differed in their effects on growth (metabolite effect: χ^2^_df=11_ = 238.75, *P* < 0.0001), and that lineages differed in their growth on individual metabolites (lineage-by-metabolite interaction: χ^2^_df=11_ = 345.16, *P* < 0.0001)(Figure 2; Table S4). The mean growth of the PAC isolates averaged across all treatments and cohorts was significantly greater than the growth of Clade D isolates (LS means: PAC = 64.4 cm^2^ x 7 days, Clade D = 31.3 cm^2^ x 7 days; *P* < 0.0001). In Clade D, growth on rutin, caffeic acid and xylose all differed significantly from the ethanol control media, while in the PAC, all metabolite treatments except chlorogenic acid, vanillic acid and starch differed significantly from the control (Table S5, Figure 2). Treatments with significant effects tended to increase growth, except caffeic acid which negatively affected the PAC (Table S6). The positive effect of caffeic acid on Clade D was such that growth rate differences between the two lineages were abolished when growing on this metabolite after correcting for multiple comparisons (LS means: PAC = 50.7 cm^2^ x 7 days, Clade D = 42.9 cm^2^ x 7 days; *P_Bonferroni_ =* 0.234). We additionally found a significant effect of experimental cohort (χ^2^_df=2_ = 126.92, *P* < 0.0001), indicating that growth rates differed between our experimental cohorts. Differences between treatments were not due to total carbon concentration, as carbon molarity was not significantly correlated with growth in either lineage across all treatments (Pearson’s r_PAC_= 0.01, *P*_PAC_= 0.83; Pearson’s r_Clade D_= 0.02, *P*_Clade D_= 0.68; Figure S1).

**Figure 2:**
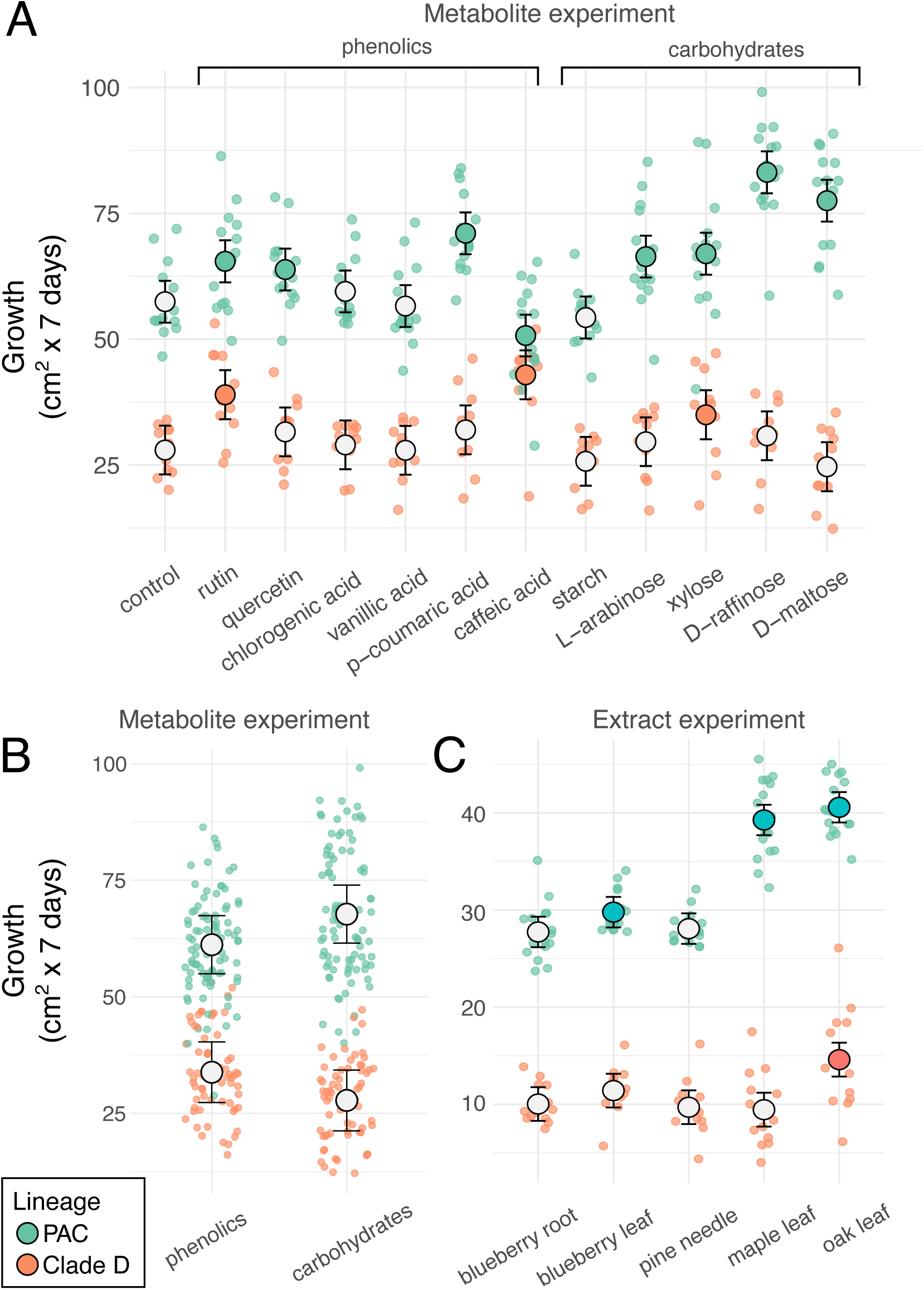
Endophyte lineages differ in their intrinsic growth rates and respond differently to specific plant metabolites, metabolite classes, and plant extracts. Dot plots displaying fungal growth (cm^2^ x 7 days), as measured by the Area Under the Growth Progress Stairs (AUGPS), in response to A) metabolite treatments, B) metabolite class and C) plant extracts. Estimated marginal means were calculated separately for the PAC and Clade D lineages and are overlaid as a large circle on top of individual data points that represent all cohorts and replicates of each isolate for each treatment. Individual data points are color coded by lineage, while treatment means that differ significantly from the control (panel A; **α** = 0.05) or from the blueberry root extract treatment (panel C; **α** = 0.05) are color coded by lineage and otherwise have a white fill if not significant. Error bars represent the 95% confidence interval, and points are arranged to be non-overlapping with jitter = 0.02.

We found that lineages differed in their baseline growth across carbohydrate and phenolic chemical classes (lineage effect: χ^2^_df=1_ = 183.38, *P <* 0.0001) and that lineages differed in their growth on individual chemical classes (lineage by chemical class interaction: χ^2^_df=1_ = 74.50, *P <* 0.0001; Figure 2B). However, phenolics and carbohydrates alone did not differ in their impact on endophyte growth (chemical effect: χ^2^_df=1_ = 0.35, *P =* 0.55; Table S7). These main effects were still significant when excluding the caffeic acid treatment (Table S8). The mean growth of PAC isolates was significantly greater than Clade D isolates on both chemical classes (Table S9); however, the magnitude of difference was greater on carbohydrates than on phenolics (LS means for carbohydrates: PAC = 67.7 cm^2^ x 7 days, Clade D = 27.8 cm^2^ x 7 days; LS means for phenolics: PAC = 61.2 cm^2^ x 7 days, Clade D = 33.8 cm^2^ x 7 days), and remained true when excluding caffeic acid treatments (Table S10). We also found a significant effect of experimental cohort (cohort effect: χ^2^_df=2_ = 32.48, *P* < 0.0001), indicating that growth rates differed between cohorts.

For the plant extract experiment, we similarly found that lineages differed in their baseline growth (lineage effect: χ^2^_df=1_ = 542.31, *P* < 0.0001), that plant extracts differed in their effect on growth (extract effect: χ^2^_df=4_ = 394.12, *P* < 0.0001), and that lineages responded differently to individual plant extracts (lineage by extract effect: χ^2^_df=4_ = 205.92, *P* < 0.0001; Table S11, Figure 2). The mean growth of PAC isolates was significantly greater than that of Clade D isolates averaged across all extract types (LS means: PAC = 33.1 cm^2^ x 7 days, Clade D = 11 cm^2^ x 7 days; *P* < 0.0001). Within the PAC, growth on the oak and maple leaf extracts was significantly greater compared with growth on all other extracts, while within Clade D, only the oak leaf treatment was significantly greater compared with all other treatments (Table S12). Although maple and oak are non-host species for these two lineages, maple and oak leaves were abundant at our study sites and may be an important resource during their saprophytic life stage.

### Lineages differ in their multivariate means across multiple metabolic traits

We calculated separate G-matrices for PAC and Clade D for both the metabolite and extract experiments using mean-centered growth data (Methods; Tables S13-20). For the metabolite experiment, the major principal component of the mean-centered G-matrices (PC1, or Gmax) for each lineage captured 73.24% and 54.71% for Clade D and PAC, respectively (Figure 3). Metabolite treatments had similar loadings to each other along each lineage’s Gmax, suggesting this axis separates isolates in both lineages according to how fast or slow they grow and independent of any specific treatment. In contrast, treatments differ in their loadings along the second principal component (PC2, capturing 7.77% and 17.42% of variation in growth rate for Clade D and PAC, respectively), which suggests that this axis separates isolates according to metabolite-specific growth responses. We confirmed a significant and large effect of endophyte lineage on mean multivariate metabolite-related growth using a one-way MANOVA (Pillai’s trace = 0.93; approximate *F* = 67.3; *df*(9,45); *P <* 0.0001; Partial eta^2^ = 0.93). For the plant extract experiment, Gmax captured 84.68% and 72.35% of variation in growth rate and PC2 captured 7.15% and 18.31% for Clade D and PAC, respectively. We observed a significant and large multivariate effect of endophyte lineage on plant extract-related mean growth using a one-way MANOVA (Pillai’s trace = 0.87; approximate *F* = 101.2; *df*(5,73); *P <* 0.0001; Partial eta^2^ = 0.87). This indicates that the lineages occupy non-overlapping regions of multivariate metabolic trait space (Figure 3B).

**Figure 3:**
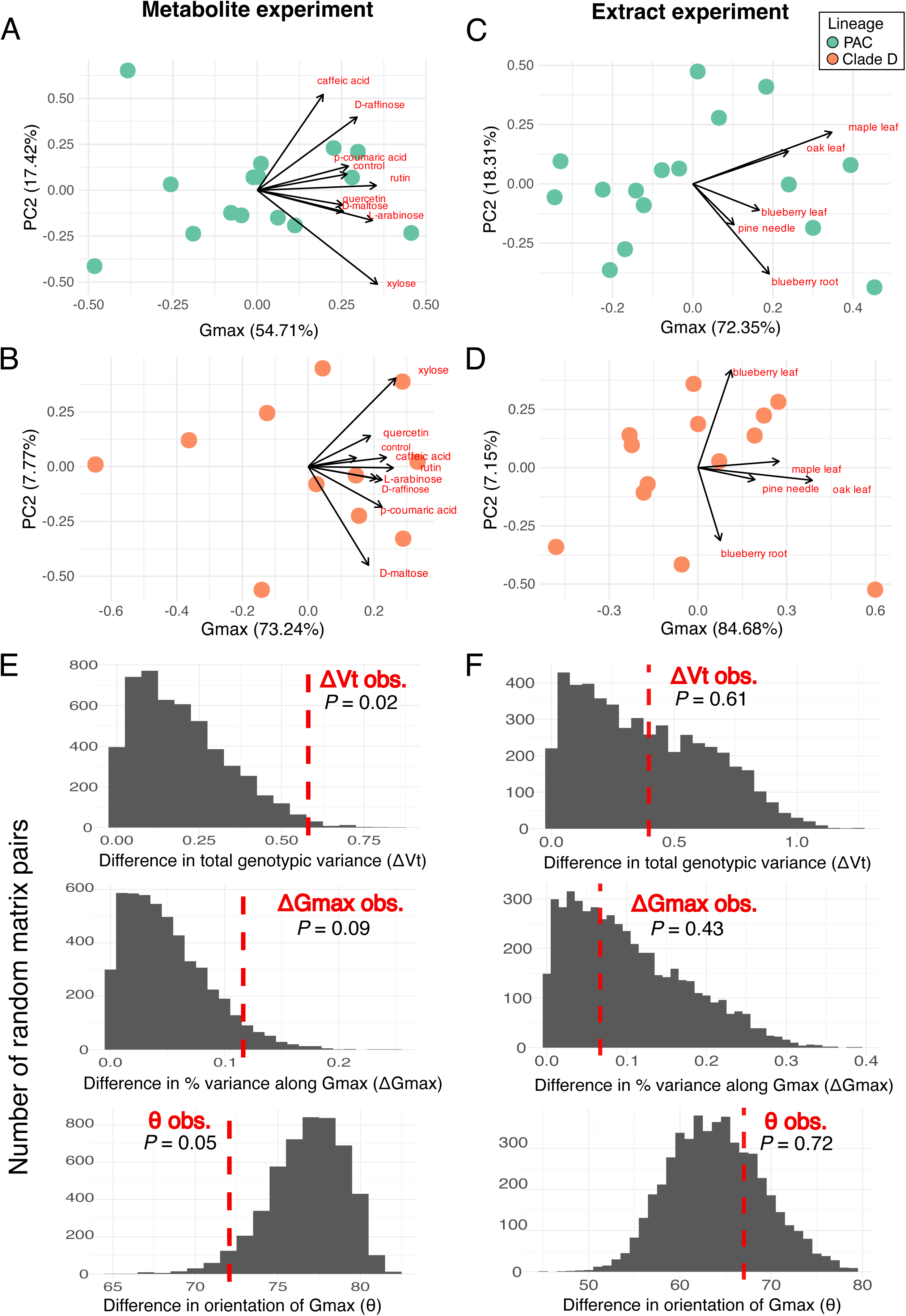
Metabolic trait architecture is largely conserved between endophyte lineages. Top: scaled biplots where the X and Y axes represent the first two principal components of the phenotypic variance/covariance (**P**) matrix for mean-centered metabolic traits in the PAC and Clade D endophyte lineages for A and B) the metabolite experiment and C and D) the extract experiment. Individual points represent individual isolates. Loadings for individual metabolites or plant extracts along PC1/Gmax and PC2 are drawn as black arrows and labeled in red. Bottom: Null distributions and observed values for the three matrix comparison statistics (difference in total phenotypic variance, difference in percent variance along PC1/Gmax and difference in the orientation of PC1/Gmax) for E) the metabolite experiment and F) the extract experiment. Null distributions were calculated by permuting isolates among lineages and comparing the features of 500 randomly permuted matrix pairs.

### Metabolic traits vary with similar strength and in similar directions along the major axis of multivariate growth

The total genotypic variance of PAC was nearly twice the magnitude of Clade D (200 vs. 110), indicating that there was more variation in growth across metabolites in PAC isolates than Clade D isolates. The difference in total genotypic variance (ΔVt) between the two lineages for the metabolite experiment was significantly greater than expected compared to random chance (*P* = 0.02; Figure 3). Lineages differed marginally in the proportion of variation explained by Gmax (ΔGmax) compared with random expectations (*P* = 0.09), indicating that the strength with which traits covary along Gmax is largely similar among isolates from either lineage. Finally, the difference in the orientation of Gmax (θ) was smaller than expected (*P =* 0.05), indicating that within each lineage, the directions in which traits covary are conserved along Gmax.

The comparisons of Gmax for the metabolite experiment, which provide comparisons along a single dimension of the G-matrix, are in agreement with comparisons of overall G-matrix structure. Random skewers showed that the response to selection predicted from the G-matrices for the two lineages was strongly correlated (random skewers correlation = 0.88, *P* <0.001). This correlation is not likely not driven by a response to a specific chemical class, as the carbohydrate-specific and phenolic-specific G-matrices were also significantly correlated across the two lineages (random skewers correlation_carbohydrates_ = 0.96, *P* = 0.005; random skewers correlation_phenolics_ = 0.88, *P* = 0.025). This indicates that overall, traits vary with similar strength and in similar directions across all dimensions of the G-matrix.

In contrast for the extract experiment, differences between lineages in total genotypic variance (ΔVt; *P* = 0.609), the proportion of variation on Gmax (ΔGmax; *P* = 0.431) and the orientation of Gmax (θ; *P* = 0.719) did not differ more or less compared to a random expectation. This indicates that PAC isolates are about as different from each other as Clade D isolates when growing on plant extracts and that their traits vary with similar strength and in similar directions (Figure 3). Similar to the metabolite experiment, a random skewers analysis showed that the G-matrices of the two lineages were positively correlated with each other (Random skewers correlation = 0.82, *P* = 0.026).

### The strength and direction of pairwise metabolic trait correlations are conserved

After comparing the overall multivariate responses of the two lineages, we next compared all pairwise trait correlations between the two lineages to test whether any substrate-specific differences in metabolic trait covariances exist. For the metabolite experiment, all trait correlations tended to be positive within each lineage, with 36/36 showing positive trends in both lineages and with 16/36 of these having positive correlations significantly greater than 0 in both lineages (Figure S2). For the extract experiment, all pairwise trait correlations within each lineage were positive, and 4/10 correlations were significantly greater than 0 in both lineages (Figure S2). After correcting for multiple comparisons, we found no evidence that pairwise trait correlations differed between lineages for the metabolite experiment, while only the oak leaf-pine needle trait correlation differed between lineages for the extract experiment (Spearman’s rho_Clade D_= 0.9, Spearman’s rho_PAC_= 0.2; Fisher’s *z* = -3.01; *P =* 0.03; Table S21, Figure 4).

**Figure 4:**
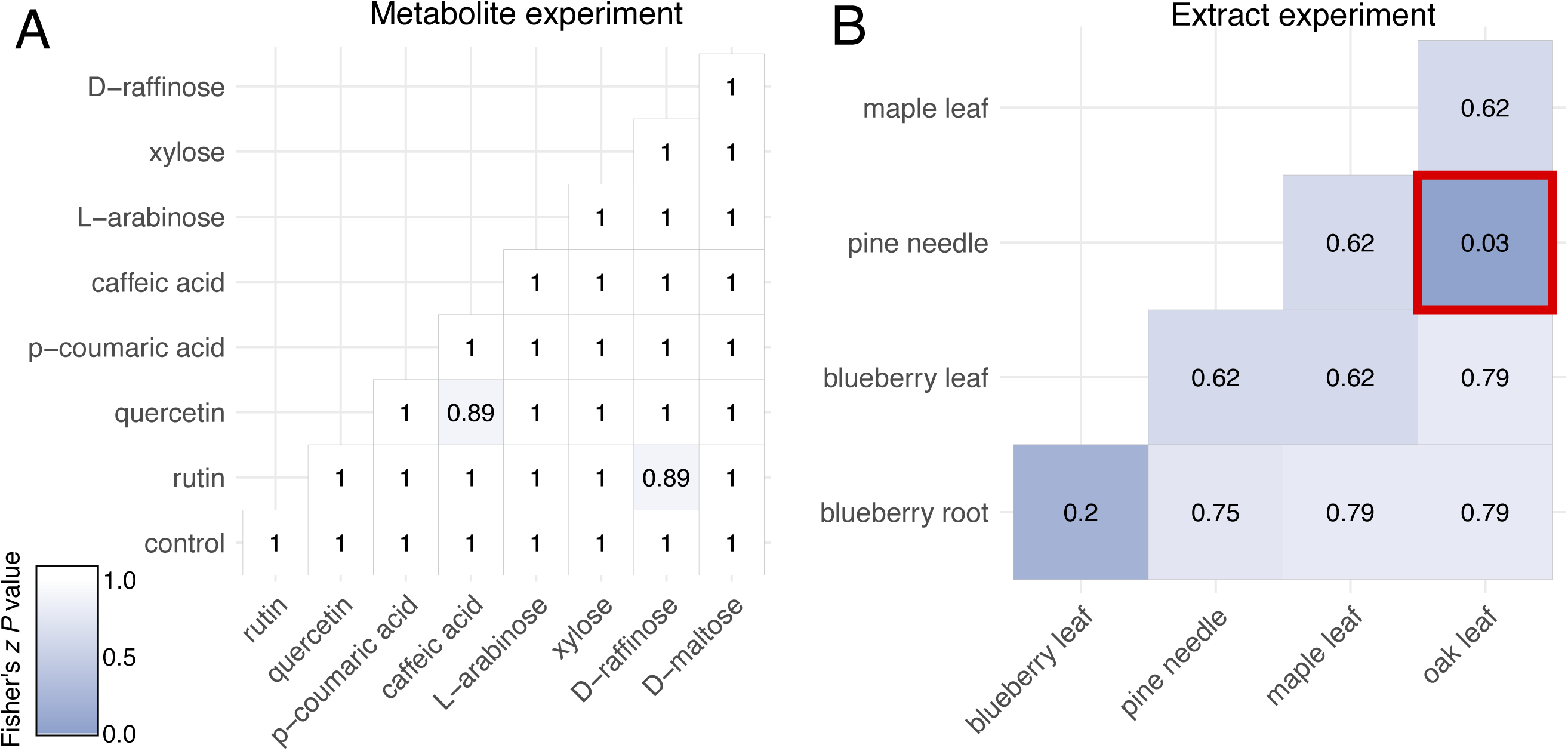
The vast majority of pairwise metabolic trait correlations do not differ between endophyte lineages. Each cell in these two matrices contains a *P* value associated with the null hypothesis that the pairwise trait correlation does not differ between the PAC and Clade D lineages for A) metabolite experiment and B) extract experiment. Statistical comparisons of Spearman’s rank correlation coefficients were conducted using the Fisher’s *z* test.

### No evidence for metabolic trait divergence along a genetic line of least resistance

We used each lineage’s G-matrix, calculated with mean-centered average growth data for each isolate, to test the hypothesis that the metabolism of lineages has diverged along a genetic line of least resistance. In contrast to previous studies that found evidence for morphological divergence along genetic lines of least resistance in multiple animal species (i.e., Schluter’s theta = 0) (Schluter 1996; McGlothlin et al. 2018), we found no evidence for metabolic divergence along a line of least resistance between the two endophytic fungal lineages in either the metabolite or extract experiment. Instead, we found that Schluter’s theta for PAC relative to Clade D and for Clade D relative to PAC for the metabolite experiment was 81.6° and 86.2°, respectively, and that theta for PAC relative to Clade D and for Clade D relative to PAC for the extract experiment was 91.3° and 83.2°, respectively.

To provide context to these results, we calculated pairwise nucleotide similarity at the ITS barcode locus between each isolate (Table S22). Mean ITS similarity was 99.6%, 96.8% and 87.2% for within PAC, within Clade D and between PAC and Clade D, respectively. A threshold of 88.5% identity at the ITS locus is typical for discriminating between taxonomic families, while species-level thresholds typically range from 97%-100% depending on the fungal taxon in question (Vu et al. 2019).

## Discussion

Here, we found that sister lineages of endophytes isolated from the same host harbor significant genotypic and phenotypic variation related to growth on plant-derived metabolites and extracts. Metabolic trait correlations were broadly conserved between lineages (Figure 3, Figure 4). However, we found no evidence of trade-offs within either lineage between specific carbohydrate and phenolic compounds, perhaps because phenolics did not inhibit endophyte growth, contrary to our initial expectation. However, lineages differed in their relative performance on phenolics versus carbohydrates, indicating that metabolic trade-offs may exist between lineages. Finally, we found that metabolic trait divergence between lineages did not occur along the genetic lines of least resistance. Divergence between the two endophyte lineages was nearly orthogonal to the major axis of genetic variation within lineages. Our results underscore the importance of species-level variation in the microbiome, and suggest that the evolutionary genetics of trait correlations in microorganisms merit further study.

### Positive correlations between metabolic traits are conserved across lineages

A central goal of our study was to test whether trait correlations are conserved across lineages in microorganisms, as they often are in macroorganisms (Arnold et al. 2008). We tested this hypothesis in two complementary experiments that measured endophyte performance across a suite of eleven host-derived metabolites and five plant extracts. In both experiments, we found that correlations between metabolic traits were largely positive and conserved between lineages (Figure 4). Neither the orientation nor magnitude of the major axis of genotypic variation differs between lineages for either metabolite- or extract-related growth phenotypes (Figure 3). Together, the conservation of pairwise positive trait correlations and the major axis of multivariate trait variation suggests that these lineages have maintained a common set of metabolic trait covariances.

Although we did not detect a difference in the magnitude or direction of genetic correlations between lineages, lineages differed in the total amount of genetic variation (ΔVt) in their response to plant metabolites (Figure 3). The PAC lineage contained almost twice the amount of genetic variation in metabolic traits compared with Clade D. This difference in the amount of standing genetic variation in the metabolic traits implies that our focal lineages differ in their evolutionary capacity to respond to selection imposed by specific metabolites (Roff et al. 2012).

The evolutionary stability of genetic (co)variances is a long-standing question in quantitative genetics (Turelli 1988; Shaw et al. 1995; Arnold et al. 2008; McGlothlin et al. 2022). All four forces of evolution—selection, drift, mutation, and gene flow—can alter genetic variances and reshape covariances between traits (Arnold et al. 2008). Indeed, the G-matrix can evolve rapidly. Selection erodes trait variance (Lande and Arnold 1983; Arnold et al. 2008), and can rapidly alter genetic correlations between traits (Steven et al. 2020). In some cases, **G** differs substantially between populations (Wood and Brodie 2015) and between species (Chakrabarty and Schielzeth 2020). Nevertheless, in many cases the structure of **G** is stable over relatively long timescales (Arnold et al. 2008, McGlothlin et al. 2022) and major environmental clines (Henry and Stinchcombe 2023). Our results are consistent with this latter group of studies, showing that the overall structure of **G** is relatively constant across lineages. We hope that our study will inspire future studies of microbial quantitative genetics, which are necessary to evaluate whether the G-matrix exhibits similar patterns of stability and evolution in micro- and macroorganisms.

We originally expected that the G-matrices in our study would be dominated by negative genetic covariances between growth on carbohydrates and phenolics. This expectation was based on the fact that sugars, fibers, and starches tend to be easily metabolized by microbes (Liu et al. 2011), while phenolics are involved in plant defense responses against microbial pathogens and tend to inhibit microbial growth (Nicholson and Hammerschmidt 1992; López-Goldar et al. 2018; Wallis and Galarneau 2020). Given the preponderance of stress-related trade-offs across taxa (Ferenci 2016; Züst and Agrawal 2017; He et al. 2022), we hypothesized that performance on growth-inhibiting compounds like phenolics would come at the expense of rapid growth on growth-promoting compounds like carbohydrates. However,we did not observe any statistically significant trade-offs between growth on these different substrates (Figure 4, Figure S2). This may be due to the fact that phenolics were not actually growth-limiting in our experiment (Figure 2A). Both Clade D and PAC grew faster on several different phenolic compounds compared with controls, suggesting that both lineages can use phenolic compounds as a nutrient source. Future studies should examine ecological traits beyond those directly related to the metabolism of plant metabolites, such as abiotic stress tolerance and combative dominance, to test if and how the metabolic traits we measured here are associated with other ecological traits (Bazzaz and Grace 1997; Crowther et al. 2014; Zanne et al. 2020).

While we did not detect a tradeoff between growth on phenolics and carbohydrates *within* lineages, comparing the lineages to each other revealed some evidence of specialization for different strategies. Clade D tended to have higher growth across all phenolics compared to all carbohydrates, while the opposite was true for PAC (Figure 2B), suggesting that Clade D in is better equipped to assimilate phenolic compounds and PAC is better equipped to assimilate carbohydrates. Indeed, we found a significant lineage x chemical class interaction effect that supports the observation that lineages respond differently to either carbohydrates or phenolics.

### Lineages diverged in baseline growth rate and in their response to specific metabolites

Although trait correlations are largely conserved between the sister lineages of Molliseaceae endophytes in our experiment, they have diverged phenotypically in two major ways. First, they differed in baseline growth rates across nearly all examined metabolites (Figure 2). The differences between lineages in intrinsic growth persisted across the plant extracts, indicating that lineages also differ in growth rate in more complex, ecologically realistic metabolic environments. In addition to differences in baseline growth, lineages differed significantly in their overall multivariate trait means on both metabolite and plant extracts (Figure 3), and in their response to specific substrates, most notably for caffeic acid (Figure 2). Together, the differences in both specific trait means and in the overall multivariate mean suggest the biochemical niches of microbes colonizing the same host are structured by both small differences across multiple traits and by large differences in specific traits.

Caffeic acid is a common metabolite produced by many plants as a precursor in the production of lignin and is also known to directly inhibit the growth of several bacteria and fungi, suggesting it has potential to act as a defense compound both through strengthening plant cell walls and inhibiting microbial colonization of plant tissues (Ravn et al. 1989; Lanoue et al. 2010; Li et al. 2021). The lineage-specific effects we observed for substrates like caffeic acid may arise through several non-mutually exclusive mechanisms. Lineages may differ in their capacity to produce enzymes that detoxify or assimilate the substrate, or may have other compensatory mechanisms for tolerating downstream stresses such as reactive oxygen species (Westrick et al. 2021).

### Divergence between lineages did not occur along the genetic line of least resistance

If trait correlations influence the trajectory of evolution, phenotypic divergence between lineages should be aligned with the multivariate trait combination containing the most genetic variation within lineages (Schluter 1996). Although trait correlations were broadly conserved between the endophyte lineages in our study, to our surprise we found that divergence in the multivariate mean metabolic phenotype was nearly orthogonal to the major axis of genetic variation within lineages. This finding is contrary to studies in macroorganisms over the past two decades that are generally consistent with Schluter’s hypothesis (Bégin and Roff 2004; McGuigan et al. 2005; Chenoweth et al. 2010; Bolstad et al. 2014; McGlothlin et al. 2018).

What might account for this pair of incongruent observations, that trait correlations are conserved between lineages, but that the major axis of genetic variation was not aligned with phenotypic divergence between lineages? One possibility is that our study omitted a major trait or traits that are correlated with the metabolic traits we measured and are under strong selection in wild endophyte populations. Mollisiaceae endophytes are facultative symbionts: they live as endophytes in blueberry roots, but are also capable of functioning as free-living saprotrophs of leaf litter (Tanney and Seifert 2020). Selection can differ dramatically between host-associated and free-living life stages in facultative symbionts (Burghardt et al. 2018). If our experiment omitted an important fitness-related trait from either life stage, indirect selection driven by the missing correlated trait could be responsible for driving evolution orthogonal to the trait correlations we measured, especially if the omitted trait or traits contain extensive genetic variation (Lande and Arnold 1983).

A second possibility is that the fungal lineages in our experiment are more distantly related than the macroorganisms that have been the predominant focus of past work. Although divergence is expected to follow the genetic line of least resistance over relatively short evolutionary timescales, this pattern may weaken as the divergence time between lineages increases, because **G** itself evolves (Schluter 1996; Arnold et al. 2008; McGlothlin et al. 2018). However, our comparisons of ITS sequences indeed suggest that the fungal lineages in our study are indeed different genera in the same taxonomic family (Tanney and Seifert 2020), similar in taxonomic scope to Schluter’s original paper, which included cross-generic comparisons in Galapagos finches and sparrows (Schluter 1996). Furthermore, a recent study in *Anolis* lizards showed that the major axis of genetic divergence shaped phenotypic divergence over relatively long timescales (20-40 million years), even in the face of an evolving G-matrix (McGlothlin et al. 2018).

Alternatively, horizontal gene transfer could account for divergence in a direction orthogonal to the genetic lines of least resistance. Once thought to be relatively rare in eukaryotes like fungi, the past two decades have revealed extensive evidence for horizontal gene transfer between fungi of genes that confer important ecological functions, including those involved in specialized metabolism (Fitzpatrick 2012; Wisecaver et al. 2014; Gluck-Thaler and Slot 2015; Szöllősi et al. 2015). The pattern we observed—divergence orthogonal to **G_max_** and little change in **G** itself—could be explained by horizontal transfer of a suite of genes that affect all measured metabolic traits. This could deliver the recipient lineage to a new region of the fitness landscape without altering trait correlations. Mobile genetic elements have previously been implicated in major evolutionary changes via horizontal transfer in bacteria, such as the transfer of antibiotic resistance or the ability to infect a host (Brockhurst et al. 2019; Weisberg and Chang 2023), and large mobile elements carrying metabolic genes and capable of horizontal transfer have recently been discovered in fungi (Vogan et al. 2021; Gluck-Thaler et al. 2022; Urquhart et al. 2022). The impact of horizontally transferable microbial mobile elements on the trajectory of evolution—the direction of evolution in multivariate trait space— in the recipient lineage, merits future research.

### Future directions: the evolutionary ecology of wild microbes

One unique challenge that arises when applying quantitative genetics to wild microbes is that the distinction between *intra*specific and *inter*specific variation is more ambiguous in microorganisms than in macroorganisms (Lücking et al. 2020). The ambiguity that arises when defining microbial species can complicate the interpretation of G-matrices. If a group of isolates is actually comprised of multiple species, the resulting G-matrix of trait correlations— which will summarize trait variation *among* species—is not a true G-matrix, and may not be a useful predictor of evolutionary trajectories *within* each constituent species. In this study, the endophyte isolates clustered into two well-resolved lineages with a high degree of sequence similarity at the ITS locus within lineages but not between lineages. This is why we considered variation within lineages to reflect intraspecific variation, and differences between lineages to reflect divergence between species. Nevertheless, the ambiguity of what constitutes a true species is a crucial factor to consider when applying quantitative genetics to microbes.

Second, it is often prohibitively difficult to control the number of isolates, lineages, or genera of wild microbes represented in a study *a priori.* It is usually infeasible to strategically target isolates or lineages for field sampling because microbial identity cannot be determined in the field. Instead, identifying wild microbes relies on culture-dependent or -independent methods back in the lab (Su et al. 2012). As a result, the number of isolates, species, or genera included in a study is often outside of the researcher’s control, at least relative to taxa that can be reliably identified and targeted for sampling in the field.

An additional complication that arises when estimating genetic (co)variances in clonal species like microbes is that some among-isolate variation may be due to transgenerational plasticity. Transgenerational plasticity is when an individual’s phenotype is determined by an environment that its parent (or grandparent or great-grandparent) experienced, rather than the alleles it inherited (Bell and Hellmann 2019). Transgenerational plasticity can inflate genetic (co)variances because it confounds genetic and environmental effects on phenotype. Accounting for this possibility in future microbial experiments requires a system-specific understanding of how rapidly these effects decay in a common garden.

Finally, the ecological significance of fine-scale taxonomic variation in host-associated microbes merits future research. Our study uncovered significant functional variation among closely related isolates in the same lineage, as well as significant functional divergence between sister lineages. How this variation scales up to influence the structure and function of host-associated communities is a major open question in the field. Through differentially impacting the growth of individual members of the plant microbiome, plant host chemistry may have cascading impacts on endophyte community assembly (Eisenhauer et al. 2017; Clocchiatti et al. 2021). Distinct host chemistries can favor the assembly of distinct endophyte communities (Sietiö et al. 2018; Bonito et al. 2019; Nickerson et al. 2023) and likely shape the distribution of endophyte populations across landscapes through environmental filtering and niche partitioning (Carroll and Petrini 1983; Tedersoo et al. 2013; Wehner et al. 2014; Veach et al. 2019). In turn, differences between endophyte communities may shape variation in plant health outcomes by, for example, influencing interactions with pathogens (Busby et al. 2013), mediating abiotic stress responses (Rodriguez et al. 2008) and promoting or inhibiting plant growth (Alberton et al. 2010; Newsham 2011; Reininger et al. 2012). Further investigations into the evolution of endophyte ecology will thus be critical for dissecting how variation within microbiomes shapes and is in turn shaped by variation in their plant hosts.

## Data availability

All supporting data, including raw image files, measurements and R scripts (including alternate scripts for analyzing ML isolates only), are available from the following Figshare repository (https://doi.org/10.6084/m9.figshare.20489715). All sequence data have been deposited at NCBI (GenBank accession numbers OP378573 through OP378602).

## Acknowledgements

We are grateful for the help of Marion Holmes and Tiffany Betras for getting us to and from our sample sites and providing expert guidance. Thanks to John Wenzel for providing access to the Powdermill Nature Reserve, Jaime Jones for facilitating our sampling at Mountain Lake Biological Station and to Vincent Formica, Chang-Yu Chang, McCall Calvert, Eunnuri Yi and Addison Martin for providing helpful feedback on drafts of this manuscript. This work was supported by startup funds from the University of Pennsylvania to CWW.

## Contributions

EGT and CWW conceptualized the research. EGT and MS collected data and EGT conducted formal analyses. EGT wrote the original draft and EGT and CWW participated in reviewing and editing the manuscript.

## Supplementary Information Legends

**Figure S1:** The linear association between carbon concentration and growth across all metabolite treatments and fungal lineages

**Figure S2:** Spearman’s rho correlations between growth across a) metabolite treatments and b) plant extract treatments for the PAC and Clade D fungal lineages. Non-significant correlations (*P* > 0.05) are crossed out.

**Table S1:** Fungal isolate ITS genotyping and best BLASTn hit from NCBI nr

**Table S2:** Carbohydrate and Phenolic metabolites known to be produced by Vaccinium species

**Table S3:** Metabolite treatments with no statistically discernible effect on the growth of either PAC or Clade D lineages

**Table S4:** Analysis of Deviance Table of the linear model with individual metabolite treatments as fixed effects for the metabolite experiment (Type III Wald χ2 tests)

**Table S5:** Estimated marginal means comparisons of all treatments vs. ethanol control in PAC and cladeD

**Table S6:** Estimated marginal means comparisons between PAC and cladeD for each metabolite treatment

**Table S7:** Analysis of Deviance Table of the linear model with carbohydrate and phenolic chemical class as fixed effects for the metabolite experiment (Type III Wald χ2 tests)

**Table S8:** Anova of fixed effects in model for phenolics vs carbohydrates, excluding caffeic acid

**Table S9:** Estimated marginal means comparisons between PAC and cladeD for carbohydrates and phenolics

**Table S10:** Estimated marginal means comparisons between PAC and cladeD for carbohydrates and phenolics, excluding caffeic acid

**Table S11:** Analysis of Deviance Table of the linear model with plant extract treatments as fixed effects for the extract experiment (Type III Wald χ2 tests)

**Table S12:** Estimated marginal means comparisons of all plant extract treatments in PAC and cladeD

**Table S13:** G-matrix for mean isolate AUGPS (Clade D, metabolite experiment, n = 11 isolates)

**Table S14:** Eigenvalues and eigenvectors of the G-matrix for mean isolate AUGPS (Clade D, metabolite experiment, n = 11 isolates)

**Table S15:** G-matrix for mean isolate AUGPS (PAC, metabolite experiment, n = 15 isolates)

**Table S16:** Eigenvalues and eigenvectors of the G-matrix for mean isolate AUGPS (PAC, metabolite experiment, n = 15 isolates)

**Table S17:** G-matrix for mean isolate AUGPS (Clade D, extract experiment, n = 13 isolates)

**Table S18:** Eigenvalues and eigenvectors of the G-matrix for mean isolate AUGPS (Clade D, extract experiment, n = 13 isolates)

**Table S19:** G-matrix for mean isolate AUGPS (PAC, extract experiment, n = 16 isolates)

**Table S20:** Eigenvalues and eigenvectors of the G-matrix for mean isolate AUGPS (PAC, extract experiment, n = 16 isolates)

**Table S21:** Tests for differences between the pairwise trait correlations of cladeD and PAC for both the metabolite and extract experiment

**Table S22:** Pairwise nucleotide identity at the ITS locus across all isolates

## References

Agrawal, A. A. 2020. A scale-dependent framework for trade-offs, syndromes, and specialization in organismal biology. Ecology 101:e02924.

Aguilar-Trigueros, C. A., S. Hempel, J. R. Powell, I. C. Anderson, J. Antonovics, J. Bergmann, T. R. Cavagnaro, et al. 2015. Branching out: Towards a trait-based understanding of fungal ecology. Fungal Biology Reviews 29:34–41.

Alberton, O., T. W. Kuyper, and R. C. Summerbell. 2010. Dark septate root endophytic fungi increase growth of Scots pine seedlings under elevated CO2 through enhanced nitrogen use efficiency. Plant and Soil 328:459–470.

Arnold, B. J., I.-T. Huang, and W. P. Hanage. 2022. Horizontal gene transfer and adaptive evolution in bacteria. Nature Reviews Microbiology 20:206–218.

Arnold, S. J., R. Bürger, P. A. Hohenlohe, B. C. Ajie, and A. G. Jones. 2008. Understanding the Evolution and Stability of the G-Matrix. Evolution 62:2451–2461.

Bazzaz, F. A., and J. Grace. 1997. Plant Resource Allocation. Elsevier.

Bégin, M., and D. A. Roff. 2004. From Micro- to Macroevolution Through Quantitative Genetic Variation: Positive Evidence from Field Crickets. Evolution 58:2287–2304.

Bell, A. M., and J. K. Hellmann. 2019. An Integrative Framework for Understanding the Mechanisms and Multigenerational Consequences of Transgenerational Plasticity. Annual review of ecology, evolution, and systematics 50:97–118.

Ben-Shachar, M. S., D. Lüdecke, and D. Makowski. 2020. effectsize: Estimation of Effect Size Indices and Standardized Parameters. Journal of Open Source Software 5:2815.

Bolstad, G. H., T. F. Hansen, C. Pélabon, M. Falahati-Anbaran, R. Pérez-Barrales, and W. S. Armbruster. 2014. Genetic constraints predict evolutionary divergence in Dalechampia blossoms. Philosophical Transactions of the Royal Society B: Biological Sciences 369:20130255.

Bonito, G., G. M. N. Benucci, K. Hameed, D. Weighill, P. Jones, K.-H. Chen, D. Jacobson, et al. 2019. Fungal-Bacterial Networks in the Populus Rhizobiome Are Impacted by Soil Properties and Host Genotype. Frontiers in Microbiology 10:481.

Boon, E., C. J. Meehan, C. Whidden, D. H.-J. Wong, M. G. Langille, and R. G. Beiko. 2014. Interactions in the microbiome: communities of organisms and communities of genes. Fems Microbiology Reviews 38:90–118.

Brockhurst, M. A., E. Harrison, J. P. J. Hall, T. Richards, A. McNally, and C. MacLean. 2019. The Ecology and Evolution of Pangenomes. Current Biology 29:R1094–R1103.

Bucknell, A. H., and M. C. McDonald. 2023. That’s no moon, it’s a Starship: Giant transposons driving fungal horizontal gene transfer. Molecular Microbiology.

Burghardt, L. T., B. Epstein, J. Guhlin, M. S. Nelson, M. R. Taylor, N. D. Young, M. J. Sadowsky, et al. 2018. Select and resequence reveals relative fitness of bacteria in symbiotic and free-living environments. Proceedings of the National Academy of Sciences 115:2425–2430.

Busby, P. E., N. Zimmerman, D. J. Weston, S. S. Jawdy, J. Houbraken, and G. Newcombe. 2013. Leaf endophytes and Populus genotype affect severity of damage from the necrotrophic leaf pathogen, Drepanopeziza populi. Ecosphere 4:art125.

Carroll, G., and O. Petrini. 1983. Patterns of Substrate Utilization by Some Fungal Endophytes from Coniferous Foliage. Mycologia 75:53–63.

Chakrabarty, A., and H. Schielzeth. 2020. Comparative analysis of the multivariate genetic architecture of morphological traits in three species of Gomphocerine grasshoppers. Heredity 124:367–382.

Chenoweth, S. F., H. D. Rundle, and M. W. Blows. 2010. The Contribution of Selection and Genetic Constraints to Phenotypic Divergence. The American Naturalist 175:186–196.

Clocchiatti, A., S. E. Hannula, M. van den Berg, M. P. J. Hundscheid, and W. de Boer. 2021. Evaluation of Phenolic Root Exudates as Stimulants of Saptrophic Fungi in the Rhizosphere. Frontiers in Microbiology 12.

Cope, O. L., K. Keefover-Ring, E. L. Kruger, and R. L. Lindroth. 2021. Growth–defense trade-offs shape population genetic composition in an iconic forest tree species. Proceedings of the National Academy of Sciences 118:e2103162118.

Crowther, T. W., D. S. Maynard, T. R. Crowther, J. Peccia, J. R. Smith, and M. A. Bradford. 2014. Untangling the fungal niche: the trait-based approach. Frontiers in Microbiology 5.

Diedenhofen, B., and J. Musch. 2015. cocor: A Comprehensive Solution for the Statistical Comparison of Correlations. PLoS ONE 10:e0121945.

Djemiel, C., P.-A. Maron, S. Terrat, S. Dequiedt, A. Cottin, and L. Ranjard. 2022. Inferring microbiota functions from taxonomic genes: a review. GigaScience 11:giab090.

Edwards, K. R., E. Kaštovská, J. Borovec, H. Šantrůčková, and T. Picek. 2018. Species effects and seasonal trends on plant efflux quantity and quality in a spruce swamp forest. Plant and Soil 426:179–196.

Eisenhauer, N., A. Lanoue, T. Strecker, S. Scheu, K. Steinauer, M. P. Thakur, and L. Mommer. 2017. Root biomass and exudates link plant diversity with soil bacterial and fungal biomass. Scientific Reports 7:44641.

Estes, S., and P. C. Phillips. 2006. Variation in pleiotropy and the mutational underpinnings of the G-matrix. Evolution 60:2655–2660.

Ferenci, T. 2016. Trade-off Mechanisms Shaping the Diversity of Bacteria. Trends in Microbiology 24:209–223.

Fitzpatrick, D. A. 2012. Horizontal gene transfer in fungi. FEMS microbiology letters 329:1–8.

Gershenzon, J., A. Fontana, M. Burow, U. Wittstock, and J. Degenhardt. 2012. Mixtures of plant secondary metabolites: metabolic origins and ecological benefits. Pages 56–77 in G. R. Iason, M. Dicke, and S. E. Hartley, eds. The Ecology of Plant Secondary Metabolites: From Genes to Global Processes, Ecological Reviews. Cambridge University Press, Cambridge.

Gluck-Thaler, E., T. Ralston, Z. Konkel, C. G. Ocampos, V. D. Ganeshan, A. E. Dorrance, T. L. Niblack, et al. 2022. Giant Starship Elements Mobilize Accessory Genes in Fungal Genomes. Molecular Biology and Evolution 39:msac109.

Gluck-Thaler, E., and J. C. Slot. 2015. Dimensions of Horizontal Gene Transfer in Eukaryotic Microbial Pathogens. PLOS Pathogens 11:e1005156.

Gompert, Z., J. P. Jahner, C. F. Scholl, J. S. Wilson, L. K. Lucas, V. Soria-Carrasco, J. A. Fordyce, et al. 2015. The evolution of novel host use is unlikely to be constrained by trade-offs or a lack of genetic variation. Molecular Ecology 24:2777–2793.

Gorzelak, M. A., S. Hambleton, and H. B. Massicotte. 2012. Community structure of ericoid mycorrhizas and root-associated fungi of Vaccinium membranaceum across an elevation gradient in the Canadian Rocky Mountains. Fungal Ecology, Fungi and Global Change 5:36–45.

Hadacek, F., and G. F. Kraus. 2002. Plant root carbohydrates affect growth behaviour of endophytic microfungi. FEMS microbiology ecology 41:161–170.

Harris, C. S., A. J. Burt, A. Saleem, P. M. Le, L. C. Martineau, P. S. Haddad, S. A. L. Bennett, et al. 2007. A single HPLC-PAD-APCI/MS method for the quantitative comparison of phenolic compounds found in leaf, stem, root and fruit extracts of Vaccinium angustifolium. Phytochemical Analysis 18:161–169.

Harrison, X. A., L. Donaldson, M. E. Correa-Cano, J. Evans, D. N. Fisher, C. E. D. Goodwin, B. S. Robinson, et al. 2018. A brief introduction to mixed effects modelling and multi-model inference in ecology. PeerJ 6:e4794.

He, Z., S. Webster, and S. Y. He. 2022. Growth–defense trade-offs in plants. Current Biology 32:R634–R639.

Henry, G. A., and J. R. Stinchcombe. 2023. G-matrix stability in clinally diverging populations of an annual weed. Evolution 77:49–62.

Ho, A., D. P. Di Lonardo, and P. L. E. Bodelier. 2017. Revisiting life strategy concepts in environmental microbial ecology. FEMS microbiology ecology 93.

Ingleby, F. C., D. J. Hosken, K. Flowers, M. F. Hawkes, S. M. Lane, J. Rapkin, C. M. House, et al. 2014. Environmental heterogeneity, multivariate sexual selection and genetic constraints on cuticular hydrocarbons in Drosophila simulans. Journal of Evolutionary Biology 27:700–713.

Katoh, K., and D. M. Standley. 2013. MAFFT Multiple Sequence Alignment Software Version 7: Improvements in Performance and Usability. Molecular Biology and Evolution 30:772–780.

Kaur, J., D. Percival, L. J. Hainstock, and J.-P. Privé. 2012. Seasonal growth dynamics and carbon allocation of the wild blueberry plant (Vaccinium angustifolium Ait.). Canadian Journal of Plant Science 92:1145–1154.

Kliebenstein, D. J., H. C. Rowe, and K. J. Denby. 2005. Secondary metabolites influence Arabidopsis/Botrytis interactions: variation in host production and pathogen sensitivity. The Plant Journal 44:25–36.

Lande, R. 1979. Quantitative Genetic Analysis of Multivariate Evolution, Applied to Brain: Body Size Allometry. Evolution 33:402–416.

Lande, R., and S. J. Arnold. 1983. The Measurement of Selection on Correlated Characters. Evolution 37:1210–1226.

Lanoue, A., V. Burlat, G. J. Henkes, I. Koch, U. Schurr, and U. S. R. Röse. 2010. De novo biosynthesis of defense root exudates in response to Fusarium attack in barley. The New Phytologist 185:577–588.

Li, S., J. Pi, H. Zhu, L. Yang, X. Zhang, and W. Ding. 2021. Caffeic Acid in Tobacco Root Exudate Defends Tobacco Plants From Infection by Ralstonia solanacearum. Frontiers in Plant Science 12.

Liu, Q., A. J. Parsons, H. Xue, K. Fraser, G. D. Ryan, J. A. Newman, and S. Rasmussen. 2011. Competition between foliar Neotyphodium lolii endophytes and mycorrhizal Glomus spp. fungi in Lolium perenne depends on resource supply and host carbohydrate content. Functional Ecology 25:910–920.

López-Goldar, X., C. Villari, P. Bonello, A. K. Borg-Karlson, D. Grivet, R. Zas, and L. Sampedro. 2018. Inducibility of Plant Secondary Metabolites in the Stem Predicts Genetic Variation in Resistance Against a Key Insect Herbivore in Maritime Pine. Frontiers in Plant Science 9.

Lücking, R., M. C. Aime, B. Robbertse, A. N. Miller, H. A. Ariyawansa, T. Aoki, G. Cardinali, et al. 2020. Unambiguous identification of fungi: where do we stand and how accurate and precise is fungal DNA barcoding? IMA Fungus 11:14.

McGlothlin, J. W., M. E. Kobiela, H. V. Wright, J. J. Kolbe, J. B. Losos, and E. D. Brodie. 2022. Conservation and Convergence of Genetic Architecture in the Adaptive Radiation of Anolis Lizards. The American Naturalist 200:E207–E220.

McGlothlin, J. W., M. E. Kobiela, H. V. Wright, D. L. Mahler, J. J. Kolbe, J. B. Losos, and E. D. Brodie. 2018. Adaptive radiation along a deeply conserved genetic line of least resistance in Anolis lizards. Evolution Letters 2:310–322.

McGuigan, K., S. F. Chenoweth, M. W. Blows, and A. E. J. M. Cheverud. 2005. Phenotypic Divergence along Lines of Genetic Variance. The American Naturalist 165:32–43.

Morvan, S., H. Meglouli, A. Lounès-Hadj Sahraoui, and M. Hijri. 2020. Into the wild blueberry (Vaccinium angustifolium) rhizosphere microbiota. Environmental Microbiology 22:3803–3822.

Nespolo, R. F., C. C. Figueroa, M. Plantegenest, and J. C. Simon. 2008. Short-term population differences in the genetic architecture of life history traits related to sexuality in an aphid species. Heredity 100:374–381.

Newsham, K. K. 2011. A meta-analysis of plant responses to dark septate root endophytes. New Phytologist 190:783–793.

Nguyen, L.-T., H. A. Schmidt, A. von Haeseler, and B. Q. Minh. 2015. IQ-TREE: A Fast and Effective Stochastic Algorithm for Estimating Maximum-Likelihood Phylogenies. Molecular Biology and Evolution 32:268–274.

Nicholson, R. L., and R. Hammerschmidt. 1992. Phenolic Compounds and Their Role in Disease Resistance. Annual Review of Phytopathology 30:369–389.

Nickerson, M. N., L. P. Moore, and J. M. U’Ren. 2023. The impact of polyphenolic compounds on the in vitro growth of oak-associated foliar endophytic and saprotrophic fungi. Fungal Ecology 62:101226.

Obeng, N., F. Bansept, M. Sieber, A. Traulsen, and H. Schulenburg. 2021. Evolution of Microbiota-Host Associations: The Microbe’s Perspective. Trends in Microbiology 29:779–787.

Parks, S. L., and N. Goldman. 2014. Maximum Likelihood Inference of Small Trees in the Presence of Long Branches. Systematic Biology 63:798–811.

Piasecka, A., N. Jedrzejczak-Rey, and P. Bednarek. 2015. Secondary metabolites in plant innate immunity: conserved function of divergent chemicals. New Phytologist 206:948–964.

Rasmussen, S., A. J. Parsons, S. Bassett, M. J. Christensen, D. E. Hume, L. J. Johnson, R. D. Johnson, et al. 2007. High nitrogen supply and carbohydrate content reduce fungal endophyte and alkaloid concentration in Lolium perenne. New Phytologist 173:787–797.

Ravn, H., C. Andary, G. Kovács, and P. Mølgaard. 1989. Caffeic acid esters as in vitro inhibitors of plant pathogenic bacteria and fungi. Biochemical Systematics and Ecology 17:175–184.

Reininger, V., C. R. Grünig, and T. N. Sieber. 2012. Host species and strain combination determine growth reduction of spruce and birch seedlings colonized by root-associated dark septate endophytes. Environmental Microbiology 14:1064–1076.

Riihinen, K., L. Jaakola, S. Kärenlampi, and A. Hohtola. 2008. Organ-specific distribution of phenolic compounds in bilberry (Vaccinium myrtillus) and “northblue” blueberry (Vaccinium corymbosum x V. angustifolium). Food Chemistry 110:156–160.

Rodriguez, R. J., J. Henson, E. Van Volkenburgh, M. Hoy, L. Wright, F. Beckwith, Y.-O. Kim, et al. 2008. Stress tolerance in plants via habitat-adapted symbiosis. The ISME journal 2:404– 416.

Roff, D. A., J. M. Prokkola, I. Krams, and M. J. Rantala. 2012. There is more than one way to skin a G matrix. Journal of Evolutionary Biology 25:1113–1126.

Rokas, A., J. H. Wisecaver, and A. L. Lind. 2018. The birth, evolution and death of metabolic gene clusters in fungi. Nature Reviews Microbiology 16:731–744.

Ruotsalainen, A. L., M. Kauppinen, P. R. Wäli, K. Saikkonen, M. Helander, and J. Tuomi. 2021. Dark septate endophytes: mutualism from by-products? Trends in Plant Science 0.

Schäfer, W., D. Straney, L. Ciuffetti, H. D. Van Etten, and O. C. Yoder. 1989. One Enzyme Makes a Fungal Pathogen, But Not a Saprophyte, Virulent on a New Host Plant. Science 246:247–249.

Schluter, D. 1996. Adaptive Radiation Along Genetic Lines of Least Resistance. Evolution 50:1766–1774.

Schneider, C. A., W. S. Rasband, and K. W. Eliceiri. 2012. NIH Image to ImageJ: 25 years of image analysis. Nature Methods 9:671–675.

Shaw, F. H., R. G. Shaw, G. S. Wilkinson, and M. Turelli. 1995. Changes in Genetic Variances and Covariances: G Whiz! Evolution 49:1260–1267.

Sietiö, O.-M., T. Tuomivirta, M. Santalahti, H. Kiheri, S. Timonen, H. Sun, H. Fritze, et al. 2018. Ericoid plant species and Pinus sylvestris shape fungal communities in their roots and surrounding soil. New Phytologist 218:738–751.

Slot, J. C., and E. Gluck-Thaler. 2019. Metabolic gene clusters, fungal diversity, and the generation of accessory functions. Current Opinion in Genetics & Development, Evolutionary genetics 58-59:17–24.

Steenwyk, J. L., T. J. B. Iii, Y. Li, X.-X. Shen, and A. Rokas. 2020. ClipKIT: A multiple sequence alignment trimming software for accurate phylogenomic inference. PLOS Biology 18:e3001007.

Ștefănescu, B.-E., L. F. Călinoiu, F. Ranga, F. Fetea, A. Mocan, D. C. Vodnar, and G. Crișan. 2020. The Chemical and Biological Profiles of Leaves from Commercial Blueberry Varieties. Plants 9:1193.

Steven, J. C., I. A. Anderson, E. D. Brodie III, and L. F. Delph. 2020. Rapid reversal of a potentially constraining genetic covariance between leaf and flower traits in Silene latifolia. Ecology and Evolution 10:569–578.

Strauss, S. Y., and R. E. Irwin. 2004. Ecological and Evolutionary Consequences of Multispecies Plant-Animal Interactions. Annual Review of Ecology, Evolution, and Systematics 35:435–466.

Stribley, D. P., and D. J. Read. 1974. The Biology of Mycorrhiza in the Ericaceae. New Phytologist 73:731–741.

Su, C., L. Lei, Y. Duan, K.-Q. Zhang, and J. Yang. 2012. Culture-independent methods for studying environmental microorganisms: methods, application, and perspective. Applied Microbiology and Biotechnology 93:993–1003.

Sun, K., F.-M. Zhang, N. Kang, J.-H. Gong, W. Zhang, Y. Chen, and C.-C. Dai. 2019. Rice carbohydrate dynamics regulate endophytic colonization of Diaporthe liquidambaris in response to external nitrogen. Fungal Ecology 39:213–224.

Szöllősi, G. J., A. A. Davín, E. Tannier, V. Daubin, and B. Boussau. 2015. Genome-scale phylogenetic analysis finds extensive gene transfer among fungi. Philosophical Transactions of the Royal Society of London. Series B, Biological Sciences 370:20140335.

Tanney, J. B., and K. A. Seifert. 2020. Mollisiaceae: An overlooked lineage of diverse endophytes. Studies in Mycology 95:293–380.

Tedersoo, L., M. Mett, T. A. Ishida, and M. Bahram. 2013. Phylogenetic relationships among host plants explain differences in fungal species richness and community composition in ectomycorrhizal symbiosis. The New Phytologist 199:822–831.

Tellenbach, C., C. R. Grünig, and T. N. Sieber. 2011. Negative effects on survival and performance of Norway spruce seedlings colonized by dark septate root endophytes are primarily isolate-dependent. Environmental Microbiology 13:2508–2517.

Turelli, M. 1988. Phenotypic Evolution, Constant Covariances, and the Maintenance of Additive Variance. Evolution 42:1342–1347.

Urquhart, A. S., N. F. Chong, Y. Yang, and A. Idnurm. 2022. A large transposable element mediates metal resistance in the fungus Paecilomyces variotii. Current Biology 32:937–950.e5.

Urquhart, A. S., A. A. Vogan, D. M. Gardiner, and A. Idnurm. 2023. Starships are active eukaryotic transposable elements mobilized by a new family of tyrosine recombinases. Proceedings of the National Academy of Sciences of the United States of America 120:e2214521120.

Veach, A. M., R. Morris, D. Z. Yip, Z. K. Yang, N. L. Engle, M. A. Cregger, T. J. Tschaplinski, et al. 2019. Rhizosphere microbiomes diverge among Populus trichocarpa plant-host genotypes and chemotypes, but it depends on soil origin. Microbiome 7:76.

Via, S. 1984. The Quantitative Genetics of Polyphagy in an Insect Herbivore. II. Genetic Correlations in Larval Performance Within and Among Host Plants. Evolution 38:896–905.

Vogan, A. A., S. L. Ament-Velásquez, E. Bastiaans, O. Wallerman, S. J. Saupe, A. Suh, and H. Johannesson. 2021. The Enterprise, a massive transposon carrying Spok meiotic drive genes. Genome Research.

Vu, D., M. Groenewald, M. de Vries, T. Gehrmann, B. Stielow, U. Eberhardt, A. Al-Hatmi, et al. 2019. Large-scale generation and analysis of filamentous fungal DNA barcodes boosts coverage for kingdom fungi and reveals thresholds for fungal species and higher taxon delimitation. Studies in Mycology 92:135–154.

Wagner, A. 2011. Genotype networks shed light on evolutionary constraints. Trends in Ecology & Evolution 26:577–584.

Wallis, C. M., and E. R.-A. Galarneau. 2020. Phenolic Compound Induction in Plant-Microbe and Plant-Insect Interactions: A Meta-Analysis. Frontiers in Plant Science 11.

Wehner, J., J. R. Powell, L. A. H. Muller, T. Caruso, S. D. Veresoglou, S. Hempel, and M. C. Rillig. 2014. Determinants of root-associated fungal communities within Asteraceae in a semi-arid grassland. Journal of Ecology 102:425–436.

Weisberg, A. J., and J. H. Chang. 2023. Mobile Genetic Element Flexibility as an Underlying Principle to Bacterial Evolution. Annual Review of Microbiology 77:null.

Westrick, N. M., D. L. Smith, and M. Kabbage. 2021. Disarming the Host: Detoxification of Plant Defense Compounds During Fungal Necrotrophy. Frontiers in Plant Science 12.

Wisecaver, J. H., J. C. Slot, and A. Rokas. 2014. The Evolution of Fungal Metabolic Pathways. PLOS Genetics 10:e1004816.

Wood, C. W., and E. D. Brodie III. 2015. Environmental effects on the structure of the G-matrix. Evolution 69:2927–2940.

Yang, H., X. Zhao, C. Liu, L. Bai, M. Zhao, and L. Li. 2018. Diversity and characteristics of colonization of root-associated fungi of Vaccinium uliginosum. Scientific Reports 8:15283.

Zanne, A. E., K. Abarenkov, M. E. Afkhami, C. A. Aguilar-Trigueros, S. Bates, J. M. Bhatnagar, P. E. Busby, et al. 2020. Fungal functional ecology: bringing a trait-based approach to plant-associated fungi. Biological Reviews 95:409–433.

Zhao, T., D. Kandasamy, P. Krokene, J. Chen, J. Gershenzon, and A. Hammerbacher. 2019. Fungal associates of the tree-killing bark beetle, Ips typographus, vary in virulence, ability to degrade conifer phenolics and influence bark beetle tunneling behavior. Fungal Ecology, From antagonism to mutualism: the chemical basis of insect-fungus interactions 38:71–79.

Züst, T., and A. A. Agrawal. 2017. Trade-Offs Between Plant Growth and Defense Against Insect Herbivory: An Emerging Mechanistic Synthesis. Annual Review of Plant Biology 68:513–534.

