## Supplementary Methods for "Multivariate divergence in wild microbes: no evidence for evolution along a genetic line of least resistance"

**Endophyte isolation protocol**

Roots from each individual plant were first washed under deionized (DI) water to remove adhering soil debris. The roots were then sanitized in a two-step process by placing them in 50ml Falcon tubes containing 30-40ml of a 4.5% active chlorine bleach: autoclaved DI water solution, vortexing for 45 seconds at max speed, transferring with sterilized forceps to another Falcon tube containing 30-40ml of a 70% ethanol: autoclaved DI water solution and vortexing again for 45 seconds. The root exteriors were then rinsed by successively transferring them to three Falcon tubes containing autoclaved DI water and vortexing for 30 seconds between each transfer. Rinsed, sanitized roots were cut into 2-5 mm fragments using sterilized scissors and placed onto 50% strength Modified Melin-Norkrans agar (MMN) plates, with approximately 5 fragments per plate and 2 plates per root sample. Plates were sealed with micropore tape and placed upside-down in dark cabinets at room temperature. Plates were checked every day for one month and any fungal mycelium that visibly emerged from root tissues was immediately sub-cultured onto fresh 50% strength MMN. To decrease the recovery of clonal isolates, we did not subculture multiple fungi with the same morphology from the same root sample. Isolates were subcultured every 7-14 days at least four times to obtain pure cultures.

**Data collection and quality control**

We calculated the Area Under the Growth Progress Stairs (AUGPS) as a quantitative measurement of growth over time on all treatments. AUGPS is conceptually similar to the Area Under the Disease Progress Curve (AUGPC), which provides a quantitative measurement of *in vivo* disease symptoms over time such that variation in growth rate among individual pathogens can be captured (Madden, Hughes, and van den Bosch 2017). AUGPS measures *in vitro* growth and is calculated by differentially weighting the first and last observations, which has been shown to improve measurement estimates over the more commonly used midpoint trapezoidal rule used to calculate AUGPC (Simko and Piepho 2012).

While measuring isolate growth, we observed unexpected morphological and growth rate variation from several replicates of the same isolate exposed to the same treatment. We thus became concerned that these cultures had become contaminated or were mislabeled. In order to identify possible mixed cultures, we calculated the growth range max(AUGPS) - min(AUGPS) for all isolate-by-treatment combinations across the metabolite experiment cohorts and the tissue experiment replicates, and identified outlier growth ranges by Tukey’s method. We removed from subsequent analysis isolates with both outlier growth ranges and whose colony morphologies differed in color and appearance across cohorts and replicates, resulting in the removal of 2 PAC isolates (ML2-1-24|27, ML1-5-39|75) and 2 Clade D isolates (ML3-6-1|49, ML1-4-61|42) from the metabolite experiment, and 1 PAC isolate (CH4-2-3|23) from the tissue experiment. We did not detect intraspecific variation in color and appearance in isolates that did not have unexpected variation in growth rate.

Furthermore, as we finalized the ITS phylogenetic tree, we lost confidence that 4 isolates had been correctly assigned to Clade D (CH3-5-11|4, CH3-2-2|88, CH3-4-2|15, CH4-5-4|35). These 4 isolates are located on a long branch whose placement within Clade D rests solely on two poorly supported branches (bootstrap support of 48 and 56). Given that Maximum Likelihood phylogenetic methods have known bias in their placement of long branches at the tips of trees (Parks and Goldman 2014), and given the poorly supported placement of these isolates within Clade D, we removed these isolates from further analysis to be conservative and decrease risk of bias when measuring lineage-specific responses. In conjunction with the filtering above, this resulted in a total of 15 PAC isolates and 11 Clade D isolates for the metabolite experiment, while Experiment had 16 PAC isolates and 13 Clade D isolates.

**DNA extraction**

We extracted DNA by collecting a small amount of aerial hyphae using a sterile pipette tip, mashing the tissue in 25μl of 1x Tris/Borate/ETDA (TBE) buffer (10.8g/L Tris base, 6.875g/L boric acid, 10ml of 0.5M EDTA) in a PCR tube, heating to 95°C for 5 minutes in a thermocycler, then micro-centrifuging at max speed for 10 seconds. We used a *Taq* DNA polymerase kit (New England Biolabs, cat.#: M0273L) with standard buffers, 10μM of the ITS1F and ITS4 primers, and 2μl of crude DNA extract for all 25μl PCR reactions. PCR amplification cycles consisted of an initial denaturing step at 95°C for 2 minutes, followed by 35 cycles of denaturing at 95°C for 30 seconds, annealing at 60°C for 1 minute, and extension at 68°C for 1 minute, followed by a final extension step at 68°C for 5 minutes. PCR products were Sanger sequenced by GeneWiz (South Plainfield, NJ, USA).

**Hot water tissue extractions**

For blueberry root extract used in the metabolite experiment, we washed roots to remove adhering soil debris, ground them into a coarse powder using an electric blender, and added 25g/L of ground tissue to flasks containing near-boiling DI water in a 95°C water bath. We incubated the flasks for 5 minutes, shook them vigorously, then incubated for an additional 5 minutes. We decanted the mixture into a clean flask through a 75μm mesh sieve, autoclaved the stock solution, and stored at 4°C overnight. The following day, we prepared root extract media by adding 40ml of the stock solution to 940ml of hot autoclaved DI water agar (15g/L agar) to obtain a final concentration of 1g root tissue/L. For plant tissue extracts used in the tissue experiment, tissues were prepared as above, except that 100g/L of ground tissue were extracted and final concentrations added to water agar media were adjusted to 5g tissue/L.

**Metabolite stock solution and media preparation**

To prepare metabolite stock solutions, we dissolved target amounts of each metabolite in pure ethanol and stored them at 4°C overnight. Using ethanol allowed us to sterilize the metabolite powder without autoclaving, since the heat and pressure from autoclaving can lead to chemical degradation. The following day, 20ml of each stock solution were mixed with warm root extract agar to obtain final target concentrations (see above). We then immediately dispensed 25ml of media into 100 x 15mm sterile Petri plates using a peristaltic pump with autoclaved tubing.
