## Supplementary figures and images for "Multivariate divergence in wild microbes: no evidence for evolution along a genetic line of least resistance"

### Figure S1

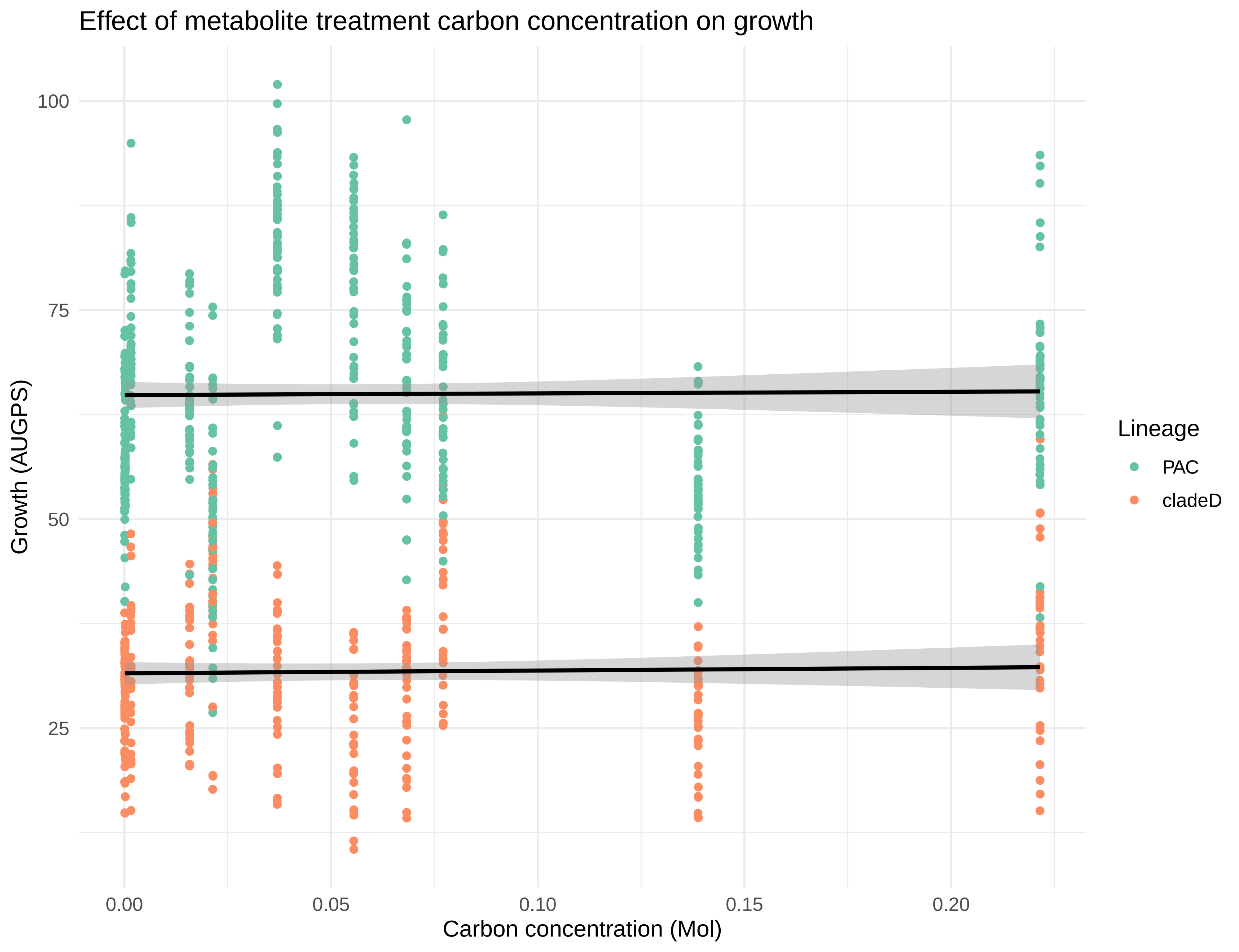

### Figure S2

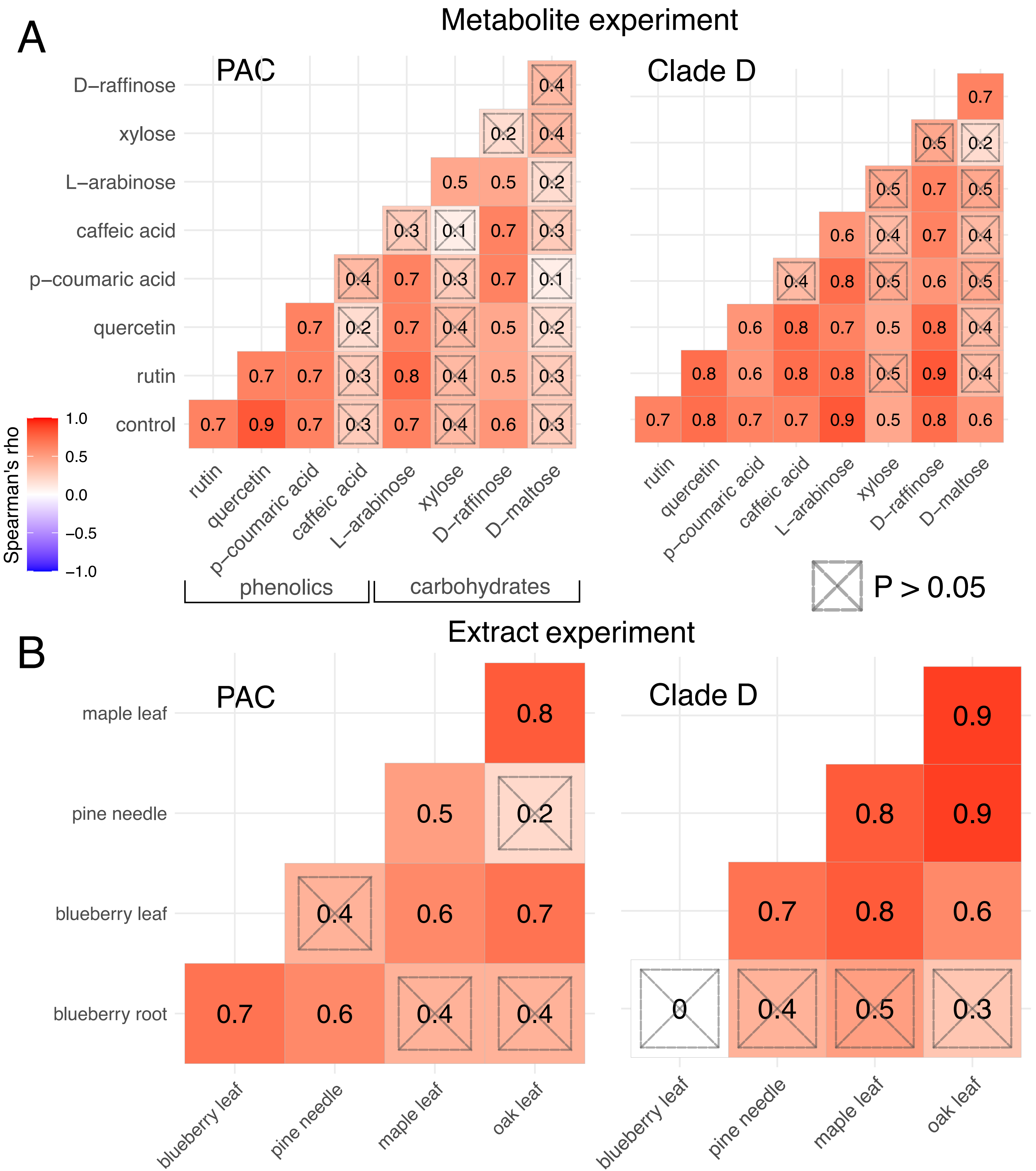
